# Hippocampal area CA2 activity supports social investigation following an acute social stress

**DOI:** 10.1101/2024.02.14.580182

**Authors:** Daniel Radzicki, Katharine E. McCann, Georgia M. Alexander, Serena M. Dudek

## Abstract

Neuronal activity in the hippocampus is critical for many types of memory acquisition and retrieval and influences an animal’s response to stress. Moreover, the molecularly distinct principal neurons of hippocampal area CA2 are required for social recognition memory and aggression in mice. To interrogate the effects of stress on CA2-dependent behaviors, we chemogenetically manipulated neuronal activity *in vivo* during an acute, socially derived stressor and tested whether memory for the defeat was influenced. One day after an acute social defeat (aSD), defeated mice spent significantly less time investigating another mouse when compared to non-defeated control mice. We found that this avoidant phenotype persisted for up to one month following a single defeat encounter. When CA2 pyramidal neuron activity was inhibited with Gi-DREADD receptors during the defeat, subject mice exhibited a significantly higher amount of social avoidance one day later when compared to defeated littermates not expressing DREADDs. Moreover, CA2 inhibition during defeat caused a reduction in submissive defense behaviors in response to aggression. *In vitro* electrophysiology and tracing experiments revealed a circuit wherein CA2 neurons connect to caudal CA1 projection neurons that, in turn, project to corticolimbic regions including the anterior cingulate cortex. Finally, socially avoidant, defeated mice exhibited significant reductions in cFos expression in caudal hippocampal and limbic brain areas during a social investigation task 24 hours after aSD. Taken together, these results indicate that CA2 neuronal activity is required to support behavioral resilience following an acute social stressor and that submissive defensive behavior during the defeat (vs. fleeing) is a predictor of future resilience to social stress. Furthermore, CA2 preferentially targets a population of caudal CA1 projection neurons that contact cortical brain regions where activity is modulated by an acute social stressor.

## Introduction

The acute stress response is conserved across species and allows an organism to learn from and adapt to a myriad of potentially threatening environmental stimuli [1–4]. Recent studies have highlighted variability in physiological and behavioral effects following exposure to a stressor. In humans, for example, a traumatic stressor can cause post-traumatic stress disorder (PTSD), which can manifest as major depressive disorder, hyperarousal, or social deficits [5, 6]. However only a subset of humans develops PTSD [7]. Similarly, only a subset of mice exposed to the chronic social defeat stress (CSDS) behavioral paradigm, in which subject mice are challenged with multiple stressful encounters with an aggressor mouse over consecutive days, develop social avoidance [8–10]. In the vulnerable, “susceptible”, mice, this chronic stress has been shown to cause long lasting changes in brain regions like the hippocampus [8, 11], striatum, and hypothalamus [12] and alter defensive behaviors in susceptible animals, while a more “resilient” population of mice exhibit minimal effects of the chronic social stressor [13]. Additionally, chronic stressors like CSDS, restraint and early-life social stress can lead to gross morphological changes in limbic regions including the prefrontal cortex and anterior cingulate [13–20]. Recent work has highlighted the transcriptomic and structural differences between vulnerable and resilient groups in an effort to better understand the mechanisms underlying stress susceptibility. For example, both the behavioral responses in a light/dark place preference assay and glucocorticoid-dependent modulation of excitatory signaling have to been shown to be predictive of which mice will go on exhibit deficits following CSDS [10, 21]. Interestingly, it is the mice that exhibit stronger defensive behaviors over repeated encounters with an aggressive mouse, that go on to show social investigative levels similar to non-defeated control mice [13].

Within the hippocampus, the morphologically and functionally distinct hippocampal area CA2 has been shown to be required for social behaviors like social recognition memory [22–27], allowing animals to form and maintain social partnerships and discern a safe familiar animal from a potential threat. This ability to discriminate between a familiar and novel conspecific, or at least show a preference for novel mice, engages downstream targets of CA2, such as caudal CA1 projection neurons, and results in activation of corticolimbic regions like the nucleus accumbens and the medial prefrontal cortex (mPFC) [28–30]. CA2 activity has also been shown to positively regulate aggression towards a conspecific via direct, excitatory projections to the dorsal lateral septum, and chemogenetic disruption of CA2 activity adversely affects context dependent fear memory in female mice [31, 32]. Furthermore, area CA2 expresses a higher density of the mineralocorticoid stress hormone receptor (MRs) when compared to the surrounding hippocampal subfields of dentate gyrus (DG), CA3 and CA1 [33, 34]. Conditionally knocking out MRs in CA2 neurons not only alters the unique molecular profile of CA2 neurons, but is also sufficient to disrupt social behavior and promote hyperarousal in response to a novel object, behaviors which are also seen in vulnerable animals following restraint stress and the CSDS paradigm [35].

Although a stressful memory may play a crucial role in an animal’s ability to predict a future threat, cognitive flexibility is also important when adapting to a changing environment. Hippocampal regions like the DG and ventral hippocampus, as well as, cortical areas like the prefrontal cortex, are required for reversal learning which, in turn, may underlie a more resilient phenotype when animals are faced with a stressor [36–38]. In fact, some research suggests that stress may facilitate this adaptive flexibility; although the underlying mechanisms are unclear, it is likely that hippocampal activity during reversal learning plays a role in the variable responses observed in a given population following either acute or chronic stress (Bryce 2015).

Given hippocampal area CA2’s established role in adaptive behaviors like social processing, aggression and contextual fear memory, and the functional role of mineralocorticoid stress hormone receptors in maintaining CA2’s molecular identity and associated behaviors, we sought to better understand the role of CA2 in stress resilience and susceptibility using an acute model of social defeat stress (aSD). Although this model has been well established in Syrian Hamsters [39, 40], recent evidence has highlighted social investigative deficits in mice following a single bout of social defeat by an aggressor [41, 42]. Using a combination of electrophysiological, immunohistochemical, behavioral, and *in vivo* chemogenetic techniques, we investigated whether the hippocampus, specifically area CA2, plays a role in social stress resilience and susceptibility in response to an acute social defeat stress. Lastly, given the lack of direct cortical projections from rostrodorsal hippocampus, we investigated how information from CA2 is routed to corticolimbic brain regions required for social processing.

## Materials and Methods

### Animals and handling

All procedures were approved by the NIEHS Animal Care and Use Committee and were in accordance with the National Institutes of Health guidelines for care and use of animals. Male and female Amigo2-icreERT2 mice were bred in house on a B6 background as described in [43] CD1 male mice were received from Charles River laboratories at 1-2 months of age. C57BL/6J (Jackson Laboratories) adult male mice (11-25 weeks of age) were used for some behavioral assays. Male and female subject mice were group housed in the same room on a 12:12 light dark cycle. CD1 male (aggressor) mice were single housed in a separate room. All behavioral assays were performed during the light cycle, between 8 AM and 3 PM. Male and female mice were used for electrophysiological and circuit tracing experiments and behavioral testing was conducted using adult male mice.

### Intracranial injections of viral vectors and tamoxifen dosing

Adult mice were temporarily anesthetized using isoflurane (0.5% at 0.8 L/min) and then received an intraperitoneal injection of a ketamine/xylazine cocktail (10 mg/ml and 0.7 mg/ml, respectively) at 0.1cc/10 g body weight. Ophthalmic ointment was then applied to the eyes and the scalp was shaved and treated with two applications of betadine. Lidocaine (0.05 cc of 0.5% Marcaine) was injected subcutaneously at the incision site. After mice were secured in the stereotaxic instrument, an incision was made along the midline exposing the skull surface and the dura was removed.

*For Gi-DREADD studies*, 500 nl of AAV5-hSyn-DIO-hM4D(Gi)-mCherry virus (Addgene #44362-AAV5) was infused bilaterally at 100 nl/min. into rostral CA2 (rCA2 :: ML: ± 2.4, AP: -2.3, DV: - 1.9) in 26 adult (18 cre- and 18 cre+, 15-19 weeks of age) Amigo2-icreERT2 mice using a Hamilton syringe positioned in a syringe pump (KD Scientific) and tube-fed cannulas made in house from blunted 26G needles (BD). The cannula was left in position for a minimum of 8 minutes before a slow retraction. The scalp was then sutured, and Buprenorphine-SR was administered subcutaneously (SQ) (0.005mg/ml, 0.02cc/10g body weight) followed by the antisedant Atipamezole (0.2mg/kg, 0.1cc dosed SQ). Animals were allowed to recover on a warming pad until regaining consciousness and returned to the home cage. Two weeks after viral infusion, Amigo2-icreERT2 mice received daily IP injections of tamoxifen (100 mg/kg dissolved in warmed corn oil) for a maximum of 7 days. Two weeks later, cre- and cre+ mice received a single dose of clozapine-n-oxide (CNO, 5 mg/kg dissolved in sterile saline) 45 minutes prior to the aSD paradigm on day 1. Following the completion of the study, expression, or lack thereof, of the mCherry tagged Gi-DREADD construct was verified in 40 µm sections from cre+ and cre- mice, respectively.

*For circuit tracing studies* involving light-evoked stimulation of rCA2 axons, 500 nl of AAV5-EF1a-DIO-ChR2(H134R)-mCherry virus (Addgene #44361-AAV5) was injected unilaterally at 100nl/min. in rostral area CA2 (ML: -2.4, AP: -2.3, DV: -1.9) in cre+ Amigo2-icreERT2 mice (N=20 mice). Two weeks later, infused mice received tamoxifen injections as outlined above. Five weeks after viral infusion, 300 µm coronal hippocampal sections were cut as described below and electrophysiological recordings conducted in caudal CA1 ipsilateral to the viral injection site.

*For retrograde tracing experiments*, 250 nl of AAVrg-CAG-mCherry virus (Addgene #59462) were intracranially injected unilaterally at 50 nl/min in the anterior cingulate cortex (ACC: ML: -0.5, AP: +0.80, DV: -1.75) of Amigo2-icreERT2 mice. A subset of Amigo2-icreERT2 mice (N=5) also received a 500 nl injection of the channelrhoposin construct (AAV5-EF1a-DIO-ChR2(H134R)-mCherry), allowing for light evoked stimulation of rCA2 axons in cCA1 cells with ACC as a known projection target.

*For Gq-DREADD mediated activation* of rostral CA2 neurons *in vivo*, 500 nl of AAV5-hSyn-DIO-hM3Dq-mCherry (Addgene #44361-AAV5) was bilaterally injected into rostral CA2 (ML: ± 2.4, AP: -2.3, DV: -1.9) in 7 adult male and female Amigo2-icreERT2 mice (4 cre- mice and 3 cre+ mice). Tamoxifen (100 mg/kg) was delivered IP two weeks later and a single dose of CNO (1 mg/kg) was given three weeks after tamoxifen. Ninety minutes following CNO, mice were transcardially perfused with 4% paraformaldehyde and the brains were sectioned for immunofluorescent staining for the immediate early gene cFos.

### Behavior and analysis

Group housed, male mice (wildtype C57BL/6J, Jackson Laboratories or GiDREADD-injected animals), 11-25 weeks of age, were placed in the home cage of adult male CD1 mice (3-10 months of age, Charles River), which had been prescreened for aggression prior to encountering the subject mice. We defined aggression as low latency (<20 seconds) to attack a novel, C57 adult male mouse over successive days. Subject mice remained in the CD1 home cage for a single, 5-minute bout of aSD and were then returned to their homecage. Twenty-four hours after the aSD, non-defeated control mice and defeated subject mice were placed in a novel arena that contained a littermate of the CD1 aggressor housed in a wire cup and allowed to explore the arena for five minutes before being returned to their homecage. In order to determine the effect of aSD over time, mice were again placed in the day 2 arena at one week, 4 weeks and, in a subset of mice, 12 weeks, that again contained a littermate of the CD1 aggressor and allowed to explore the context for five minutes. Video recordings were taken during both the defeat and the investigative encounters for behavioral analysis. Defeat videos were analyzed by an experienced individual, blinded to genotype as applicable, and the following behaviors were scored: amount of direct aggression a subject mouse received (seconds); passive submission during aggression + subject flee reactions when facing aggression (seconds); defensive submission, defined as a subject mouse taking a defensive, often upright posture and standing its ground when facing aggression (seconds). Passive and defensive submission scores were later added together in order to calculate total submission during defeat. For day 2 videos, the novel area was divided in half with the half furthest from the contained CD1 mouse defined as the “far zone” and the immediate area (16 cm diameter) surrounding the wire cup defined as the “interaction zone” (**Figure1a**). EthoVision video tracking software (Noldus) was used to score the following subject behaviors during the day 2 avoidance/investigation assay; total distance travelled (cm), time spent in the far zone (seconds), time spent in the interaction zone (duration, seconds), and number of distinct entries in the interaction zone. Avoidant mice were defined as mice that spent more than 200 seconds of the 300 second screen in the far zone of the arena. Data are presented as mean ± SEM.

A subset of adult male C57 mice (N=11) underwent a two-day fear conditioning protocol. During day one fear acquisition, mice were individually housed in fear conditioning chambers (Med Associates) for a three-minute habituation period followed by five tone (75db)/shock (0.6) pairings separated by three-minute intervals. Freezing behavior during the habituation period and five post-shock intervals was calculated using the Video Freeze program. On day 2, ∼24 hours after the fear acquisition task, mice were returned to the day 1 context and freezing behavior was again monitored and calculated using Video Freeze. The chambers were cleaned with 75% isopropyl alcohol and five sprays of Simple Green cleaning solution between every trial [44].

Five weeks following a CA2 targeted, intracranial injection of the Gi-DREADD construct (and 3 weeks following the first of the IP injections of tamoxifen) Amigo2-icreERT2 mice received an IP injection of clozapine n-oxide (CNO) at 5 mg/kg 45 minutes prior to or immediately following a 5-minute bout of aSD and then tested for avoidance/investigation 24 hours later. Following the day 2 assay, a subset of non-defeated and defeated Amigo2-icreERT2 mice underwent cardiac perfusion with a 5% paraformaldehyde (PFA) solution 45 minutes after the assay in order to preserve the brains for immunostaining.

#### Patch clamp recordings in acute hippocampal slices

Adult Amigo2-icreERT2 mice (N=20, male and female) received an IP dose of Fatal Plus (sodium pentobarbital, 50 mg/mL, ≥100 mg/kg) and were transcardially perfused with 10 ml of ice-cold ACSF solution containing the following (in mM): 125 NaCl, 2.5 KCl, 1.25 NaH_2_PO_4_, 26 NaHCO_3_, 10 glucose, 2 Na-pyruvate, 0.5 CaCl_2_, and 7 MgCl_2_, 100 picrotoxin (Abcam), saturated with 95% O_2_ and 5% CO_2_ per a protocol developed for recording from adult mouse slices [45]. The brain was removed from the skull in ice-cold ACSF solution. Coronal slices at 300 µm thickness were cut using a vibratome (Leica VT-1200), stored for ∼60 min at 32°C, and allowed to recover at room temperature (22– 24°C) for at least 30 min in ACSF (saturated with 95% O_2_ and 5% CO_2_). For recordings, slices were transferred to a recording chamber and continuously superfused with ACSF containing the following (in mM): 125 NaCl, 2.5 KCl, 1.25 NaH_2_PO_4_, 26 NaHCO_3_, 25 glucose, 2 CaCl_2_, and 1 MgCl_2_, saturated with 95% O_2_ and 5% CO_2_. All recordings were performed using an Axopatch 200B amplifier. Signals were filtered at 10 kHz and sampled at 20 kHz for current-clamp recordings and filtered at 5 kHz and sampled at 10 kHz for voltage-clamp recordings. Pipettes were pulled from WPI glass (PG10 –165) using a horizontal puller (P97; Sutter Instruments) and were filled with an internal solution consisting of 140 K-gluconate, 10 HEPES, 0.1 EGTA, 8 NaCl, 4 MgCl_2_, 2 ATP, and 0.3 GTP, along with biocytin (1 mg/ml), pH 7.3 with KOH. Pipette resistances in the working solutions ranged from 3 to 6 MΩ resulting in series resistances of 10-30 MΩ. rCA2 axons were stimulated using 1-ms blue light pulses generated by a 455 nm LED (Prizmatix UHP-T-455-EP) mounted on a Zeiss Examiner Z.1 microscope and directed through a Zeiss 40x objective.

### Tissue processing and immunohistochemistry

Following the conclusion of a given experiment, mice were deeply anesthetized with Fatal Plus (sodium pentobarbital, 50 mg/mL: >100 mg/kg, IP). Immediately after transcardial perfusion with 4% paraformaldehyde (pH 7.4) in phosphate buffered saline (PBS), brains were removed and post-fixed in the same fixative overnight at 4°C. Brains were coronally sectioned at 40 µm on a Leica VT 1200S vibratome. Brain sections were then stored free floating in PBS with 0.02% sodium azide at 4°C until staining.

Unless otherwise noted, all free-floating sections were washed in 1xPBS for 15 minutes and incubated overnight at 4⁰C in 3% neuronal goat serum (NGS) in 0.3% PBS-Triton followed by a 2-day incubation in primary antibody in 3% NGS/0.3% PBS-Triton at 4⁰C. Sections were then washed for 20 minutes in 0.3% PBS-Triton and then 3 times 0.3% PBS-Triton for 10 minutes each before being incubated overnight at 4⁰C in secondary antibody in 3% NGS/0.3% PBS-Triton. For identification of biocytin filled CA1 neurons in the caudal hippocampus, sections were stained with a fluorescence-tagged streptavidin secondary antibody. Finally, sections were washed once in 0.3% PBS-Triton for 30 minutes and three time in 1xPBS for 15 minutes each at room temp before being cover slipped with Vectashield Hardset Mounting Medium with DAPI (Vector, H-1500-10).

The following antibodies were used to stain CA2 axons (1⁰ chicken anti-GFP (Abcam #ab13970) 1:1000 + 2⁰ anti-chicken 488 (Invitrogen #A11039) 1:500) and biocytin filled cells (2⁰ Streptavidin-568 (Invitrogen #S11226) 1:500) in caudal hippocampus. A subset of sections was stained for calbindin+ cells (1⁰ rabbit anti-calbindin (Swant #CB38) 1:500 + 2⁰ anti-rabbit-633 (Invitrogen #A21071) 1:500). To visualize Gi-DREADD-mCherry expression in rCA2 of injected mice sections were stained with 1⁰ rat anti-mCherry (Life Technology #M11217) 1:500 + 2⁰ anti-rat 568 (Invitrogen #A11077) at 1:250. In order to visualize AAV-retro positive cells sections were stained for mCherry (1⁰ rat anti-mCherry (Life Technology #M11217) 1:500 + 2⁰ anti-rat 568 (Invitrogen #A11077) at 1:250) paired with (2⁰ Streptavidin-633 (Invitrogen #S21375) 1:500).

For immunofluorescent staining of the IEG cFOS, free-floating sections were washed twice in 1x PBS for five minutes followed by a permeabilization wash in 0.3% PBS-Triton for 20 min and a 5-minute wash in 1x PBS. Cells were incubated at room temperature (RT) in 10% normal goat serum (NGS) in 0.1% PBS-Triton for 1 hour. Sections were then incubated overnight at 4⁰C on a plate rocker in a 5% NGS/0.1% PBS-Triton solution containing rabbit anti-cFos antibody (Abcam, ab190289) at 1:500 dilution. On day 2 sections were allowed to incubate at RT for ∼2 hours before 2 5-minute washes in 0.1% PBS-Triton, a third 5-minute wash in 1xPBS and 2 hours in Goat anti-rabbit 488 (Thermo A11034) at a 1:500 dilution in 5% NGS/0.1% PBS-Triton at RT. Following secondary incubation, sections were washed twice in 0.1% PBS-Triton for 10 minutes, and once in PBS for a minimum of 20 minutes. Lastly, sections were given a final 5-minute wash in 1xPBS, and cover slipped with Vectashield Hardset Mounting Medium with DAPI (Vector, H-1500-10).

### Microscopy and analysis

All images were acquired with an LSM 880 Zeiss confocal microscope. For images that were later used for quantification, imaging parameters were kept constant for sections of a given brain region. All quantifications were made on maximum intensity *z*-projected images with ImageJ [46]. Data are presented as mean ± SEM.

#### Determining the location of a biocytin filled caudal hippocampal neuron along the dorsal-ventral axis

Following the completion of a whole-cell electrophysiological recording, the recording pipette was slowly retracted allowing the soma to reseal in order to preserve the biocytin labelling. Sections were then transferred to 4% PFA overnight at 4⁰C before being transferred to PBS. Next, sections were immunostained with Streptavidin-568 (Invitrogen #S11226) and, in a subset of sections, calbindin (1⁰ rabbit anti-calbindin (Swant #CB38) + 2⁰ anti-rabbit-633 (Invitrogen #A21071)). Prior to imaging, the 400 µm sections were incubated at room temperature in a thiodiethanol solution (30% in PBS, Sigma-Aldrich #88561) for a minimum of 30 minutes rendering the sections translucent. Confocal images of the caudal hippocampus were taken and the distance from the dorsal to the ventral end of the CA1 cell body layer was measured using Image J. This measurement was then divided into three equal parts in order to discriminate between the dorsal, intermediate, and ventral subregions of caudal CA1. Imaged biocytin labelled neurons were then categorized as residing in one of these three subregions.

#### Quantification of cFos+ cells following the social avoidance assay

Sections containing the hippocampus, lateral septum and frontal cortical brain regions used in the quantification of cFos+ cells were immunostained using the protocol detailed above. 20x tile scan images of our regions of interest were taken at 512×512 resolution at a *z*-section depth of 4.04 µm, 4-5 *z*-sections per stack. Laser power, pinhole size, and exposure settings were kept consistent for a given brain region and set to a level that maximized the fluorescent signal while minimizing background for a given region. Prior to quantification 3 *z*-sections were max projected for a total quantifiable depth of 12 µm per section. Regions of interest (ROIs) were drawn in Image J to further subdivide brain regions to be quantified based on landmarks and measures from The Mouse Brain in Stereotaxic Coordinated, 2^nd^ edition [47]. cFos+ cell counts were achieved using the Image J Image Processing analysis plugin. Grayscale images were first thresholded consistently across sections for a given brain region and particles (“Analyze Particles”) were analyzed with the following constraints: size µm^2^=400-1000, circularity=0.5-1.0. A minimum of 2 hemispheres were quantified for a given brain region and cell counts were averaged per animal. cFos quantification data presented as mean ± SEM.

## Results

### Acute social defeat results in long lasting social avoidance

Repeated, stressful exposure to an aggressive mouse over multiple days results in behavioral and physiological changes in subject animals, but less is known about how an acute aggressive defeat alters mouse investigative behavior. To better understand these potential effects, we subjected adult male, wildtype mice (C57BL/6J) to a 5-minute bout of aSD in the home cage of a male CD1 aggressor mouse. One day after this stressful encounter, subject mice were placed in a novel arena that contained a male littermate of the CD1 aggressor and were allowed to explore the context for 5 minutes. We then calculated the amount time a subject mouse spent in the far zone of the area (avoidance) and the number of discrete entries into the interaction zone surrounding the CD1 mouse (investigation) (**Figure 1a**). When compared to non-defeated control mice, defeated mice spent significantly more time in the far zone of the arena (**Figure 1b,c,d**). Additionally, they exhibited significantly fewer entries into the interaction zone (**Figure 1e**). Not only did aSD result in a reduction in social investigation, but defeated mice also covered significantly less total distance over the course of the 5-minute avoidance assay (**Suppl Figure 1c**). We next asked if any metrics observed during the day 1 defeat would predict the degree of social avoidance a defeated subject mouse would exhibit 24 hours later. We observed no significant correlation between either the amount of aggression a subject received and the amount of time of spent in the far zone (**Supplementary Figure 1a**) or total observable submissive behaviors such as freezing, flee, and defensive posturing and avoidance 24 hours post defeat (**Supplementary Figure 1b**). Surprisingly, this avoidant phenotype persisted in the more “susceptible” population of defeated mice: when re-exposed to the day 2 context, containing a novel CD1 mouse, one month after the initial acute defeat exposure, defeated mice that spent >200 seconds in the far zone of the arena on day 2 (**Figure1f,** red) continued to exhibit significantly higher levels of avoidance when compared to more investigative cohort of defeated mice (**Figure1f**, blue, **Supplementary Figure 1d**). Moreover, this avoidant phenotype is observable up to 3 months in the most susceptible mice following a single bout of social aggression (**Supplementary Figure 1d**).

**Figure 1:**
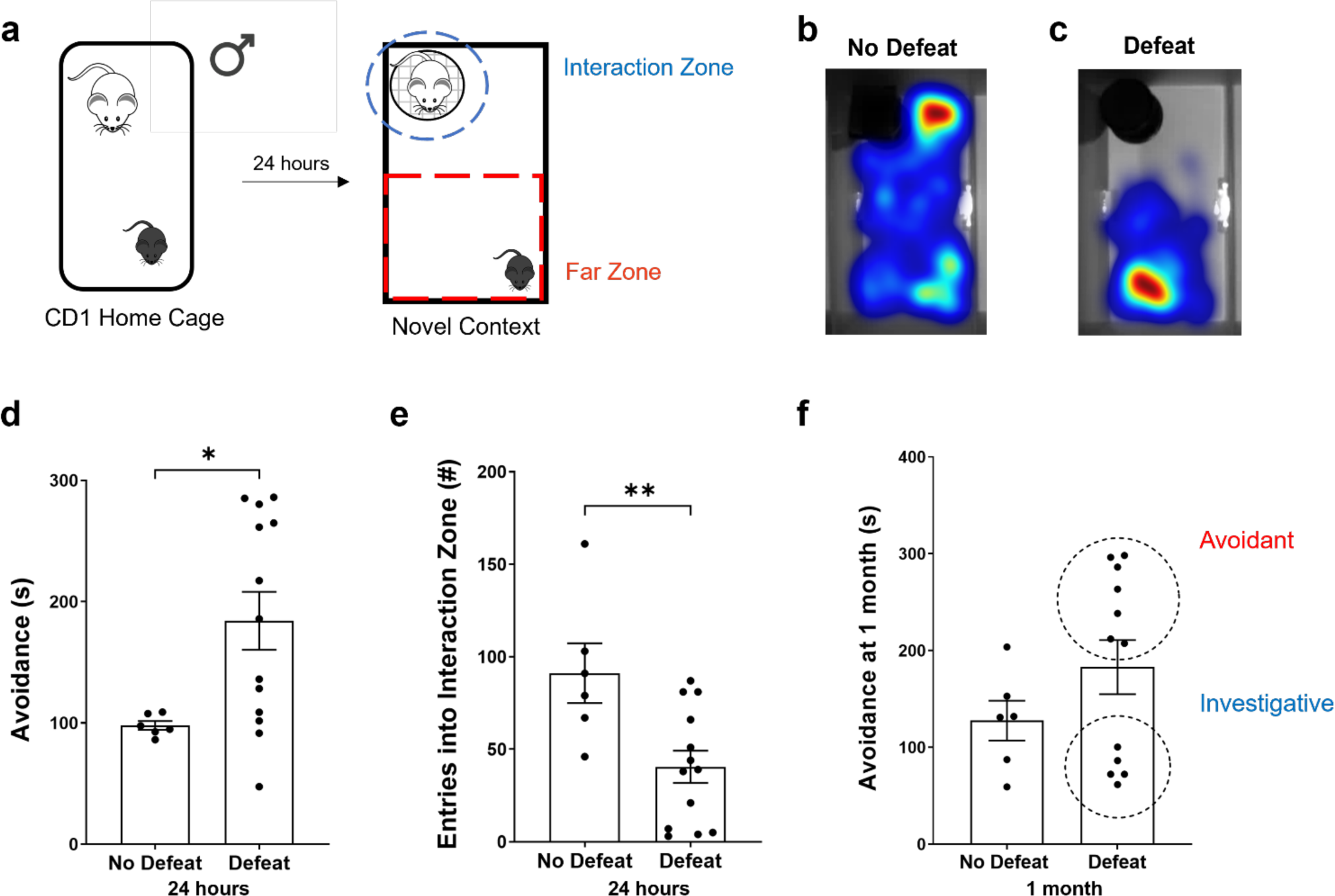
Acute social defeat results in social avoidance of a novel mouse. **a.** In the acute social defeat model (aSD) a male subject mouse (C57BL/6J, 11-25 weeks of age) was placed in the home cage of an adult male CD1 for a five-minute bout of defeat then returned to the home cage. Twenty-four hours later the subject mouse was allowed five-minutes to explore a novel environment containing a littermate of the CD1 aggressor housed within a wire cup. Warm colors represent more time spent. **b.** Heatmap illustrating a non-defeated subject’s preference investigation of the novel CD1 mouse. **c.** Heatmap illustrating a defeated subject’s preference for the “far zone” opposite the location of the CD1. **d.** Defeated mice spent significantly more time in the “far zone” of the arena when compared to non-defeat mice 24 hours after aSD (No Defeat: 97.98 ± 3.6 seconds vs. Defeat: 190.49 ± 25.0 seconds, N=6 & 12 mice, respectively, unpaired *t-test p*=0.02). **e.** Non-defeated control mice interacted with the CD1 mouse at a significantly higher frequency, measured as discrete entries into the interaction zone, than the defeated mice 24 hours after aSD (No Defeat: 91.17 ± 16.1 entries vs. Defeat: 40.25 ± 9.4 entries, N=6 & 12 mice, respectively, unpaired *t-test p*=0.01). **f.** One month following a single bout of defeat the two groups, as a whole, no longer exhibited significantly different measures of avoidance (No Defeat: 127.80 ± 20.6 seconds vs. Defeat: 180.75 ± 30.5 seconds, N=6 & 11 mice, respectively, unpaired *t-test p*=0.25). Notably, though, a more avoidant subpopulation (>200 seconds spent in the far zone) of defeated mice emerged as distinct from the more investigative mice (Defeat-Avoidant: 265.81 ± 14.2 seconds vs. Defeat-Investigative: 78.68 ± 6.7 seconds, N=6 & 5 mice, respectively, unpaired *t-test p*<0.0001). * *p<*0.05, ** *p*<0.01.

### Chemogenetic inactivation of CA2 pyramidal neurons increases avoidance after an acute social defeat

Given the now well-established role of hippocampal area CA2 in behaviors such as social discrimination memory and aggression, we next asked whether inhibition of neuronal activity in this region during aSD would alter the avoidant/investigative phenotype 24 hours following defeat. To that end, we injected intracranially a cre-dependent inhibitory Gi-DREADD AAV into area CA2 of cre+ and cre-Amigo2-icre-ERT2 male mice. Forty-five minutes prior to the defeat, CNO was delivered intraperitoneally to both populations of subject mice, effectively silencing CA2 neuronal output in subject mice (cre+ defeat group, expressing Gi-DREADD) during the stressful defeat encounter [32]. Cre- and cre+ mice were then subjected to the day 2 avoidance test as previously outlined (**Figure 2a,b**). Similar to what we observed in male C57 wildtype, defeated cre- mice (no DREADD-expressing control mice), with CA2 activity left unperturbed, spent significantly more time in the far zone of the arena (**Figure 2c**) with significantly fewer entries into the interaction zone surrounding the CD1 mouse (**Figure 2d**) than non-defeated control mice. Surprisingly, cre+ mice, with CA2 neuronal activity inhibited during defeat, exhibited an even stronger avoidance phenotype when compared to cre-defeat counterparts as measured by time spent in zone opposite the CD1 mouse (**Figure 2c,d**) and number of entries in the surrounding interaction zone (**Figure 2c,d**). The significant increase in social avoidance in mice where CA2 activity was inhibited during the defeat was driven by the loss of the more investigative/”resilient” population of mice that compose ∼50% of the cre-defeat group (**Figure 2e**). As CNO-driven activation of an inhibitory Gi-DREADD is known to exert its effects for up 6 hours following administration, we next wondered if the investigative deficits we observed in cre+ mice were due to CA2 neuronal inhibition during acquisition or a result of Gi-DREADD mediated disruption of encoding or consolidation of the stress memory which occurs on the order of hours. To better understand the role of these distinct phases of memory processing we ran a second cohort of Gi-DREADD expressing mice through the aSD paradigm with CNO being delivered to both cre- and cre+ defeat groups immediately following the defeat (**Supplementary Figure 2a**). Chemogenetic inactivation of area CA2 immediately following the defeat encounter had no statistically significant effect on avoidance levels (**Supplementary Figure 2b**), whereas both groups showed significantly less avoidance levels relative to defeated cre+ animals that had CA2 activity inhibited during defeat (**Supplementary Figure 2b**).

**Figure 2:**
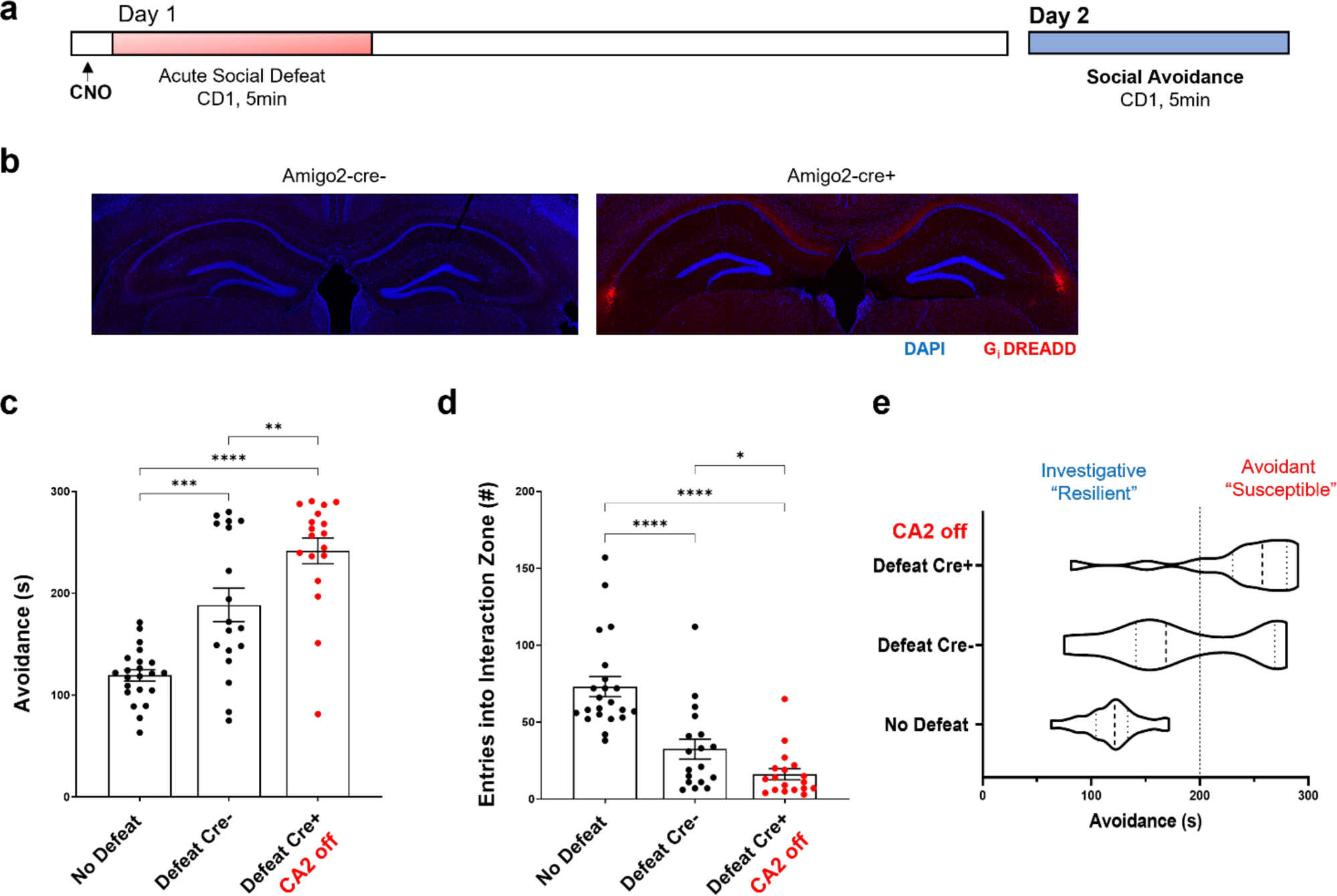
Chemogenetic inactivation of CA2 alters investigation of a CD1 mouse 24 hours after defeat. **a.** Acute social defeat experimental timeline. Five mg/kg of clozapine-N-oxide (CNO) was delivered on day 1, 45 minutes prior to defeat by a CD1 aggressor mouse via IP injection in Amigo2-icreERT2 male mice (15-19wks of age) that either did (Cre+) or did not (Cre-) express cre-recombinase in area CA2 neurons. Twenty-four hours after defeat, day 2, mice were tested for avoidance/investigation in a novel context containing a littermate of the CD1 aggressor mouse. **b.** (right) Amigo2-icreERT2-Cre+ mice show CA2 specific expression of the intracranially injected Gi-DREADD-mCherry construct 3+ weeks following injection (left). No expression of the Gi-DREADD construct was observed in cre- mice. **c.** Day 2, 24 hours after defeat, both defeat populations spent significantly more time in the far zone of the novel arena, opposite the CD1 mouse (i.e. avoidance behavior), when compared to no defeat control mice. Cre+ defeated males, with CA2 activity inhibited, spent significantly more time in the far zone than Cre-defeated mice (No Defeat controls: 119.44 ± 5.7 seconds vs. Defeat Cre-: 188.60 ± 16.4 seconds vs. Defeat Cre+: 241.64 ± 12.7 seconds, N=22, 18, 18, respectively, one-way ANOVA F=28.19 *p*<0.0001; No Defeat vs. Defeat Cre-, Holm-Sidak multiple comparisons test *p*= 0.0.0002, No Defeat vs. Defeat Cre+, Holm-Sidak multiple comparisons test *p*<0.0001, Defeat Cre-vs. Defeat Cre+, Holm-Sidak multiple comparisons test *p*=0.003). **d.** Day 2, 24 hours after defeat, Cre+ mice entered the interaction zone, surrounding the CD1 mouse, at a significantly lower frequency than both Defeat Cre- and No Defeat control mice (No Defeat controls: 73.09 ± 6.5 entries vs. Defeat Cre-: 32.5 ± 6.5 entries vs. Defeat Cre+: 16.17 ± 3.4 entries, N=22, 18, 18 mice, respectively, one-way ANOVA F=26.34 *p*<0.0001; No Defeat vs. Defeat Cre-, Holm-Sidak *p*<0.0001, No Defeat vs. Defeat Cre+, Holm-Sidak p<0.0001, Defeat Cre- vs. Defeat Cre+, Holm-Sidak *p*=0.062, unpaired *t-test p*=0.034). **e.** The increase in avoidance behavior (red) in Defeat Cre+ mice was driven by a loss of the more investigative phenotype (green) when CA2 activity was inhibited during aSD. * *p<*0.05, ** *p*<0.01, *** *p*<0.001, **** *p*<0.0001.

### Chemogenetic inactivation of CA2 alters defensive behaviors during defeat

A single bout of received aggression is sufficient to drive social avoidance in a subset of defeated mice. Given our observation that CA2 activity during defeat supports later investigative behaviors, we next sought to determine whether a subject mouse’s behavior during the defeat is predictive of future investigation/”resilience” or avoidance/”susceptibility”. On day 1 of the aSD paradigm with CNO delivered 45 minutes prior to defeat (**Figure 3a**), we observed no significant correlation between the amount of aggression received and future avoidance in cre- and cre+ defeated mice (**Figure 3b**). Similarly, we observed no significant correlation between the amount of total submissive behaviors expressed by mice during the defeat encounter and time spent in the far zone on day 2 of the paradigm (**Figure 3c**: (N=36 mice, R^2^=0.0297, *p*>0.005). Because mice are capable of a range of submissive behaviors when faced with threat or aggression, we next subdivided total submission into either submissive flee (time spent freezing or fleeing the aggression) or a more defensive response where the subject would raise up on their hindlegs and stand ground in response to CD1 aggression (defensive submission). Mice that showed high levels of submissive flee during defeat went on to display higher measures of avoidance on day 2 as evidenced by a positive correlation between these two measures (**Figure 3d**), though we found no significant difference in absolute amount of submissive flee between defeated cre- and cre+ mice (**Figure 3e**). When CA2 was inhibited during the defeat, cre+ mice spent significantly less time in defensive postures (**Figure 3f**) and showed a significant reduction in defensive submission as a percentage of total submissive behaviors (**Figure 3g**). Confirming that this cohort of mice receiving CNO immediately following the aggressive encounter were similar to each other during the defeat, we observed no difference in behavior between cre- and cre+ mice in response to the aggressor (**Supplementary Figure 2c**).

**Figure 3:**
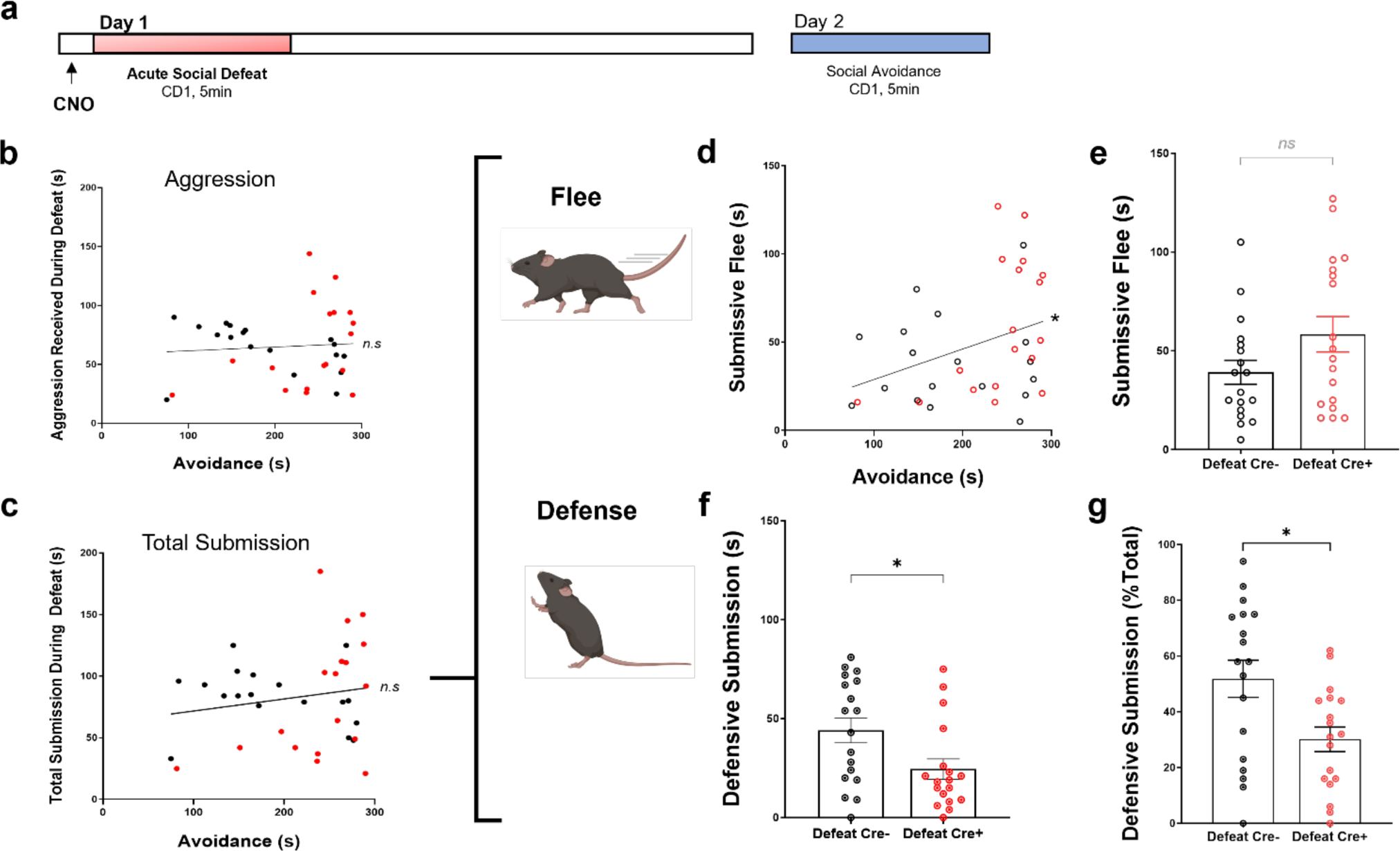
Chemogenetic inactivation of CA2 alters subject behavior during defeat. **a.** Acute social defeat experimental timeline. **b.** No significant correlation was observed between the amount of aggression a subject mouse received during the day 1 defeat and the amount of time spent in the “far zone” of the arena (i.e. avoidance) 24 hours after defeat (N=36 mice, R^2^=0.0052, *p*>0.005). **c.** No significant correlation was observed between the amount of total submission (combined flee/freezing + defensive behaviors) exhibited by subject mice during defeat and avoidance 24 hours following the defeat (N=35 mice, R^2^=0.0297, *p*>0.005). **d.** During the 5-minute defeat, the amount of submissive flee/freezing behaviors exhibited by subject mice positively correlates with an increase in CD1 avoidance 24 hours after defeat (N=36 mice, R^2^=0.1196, *p*=0.039). **e.** The absolute amount of submissive fleeing/freezing did not significantly differ between the Cre- and Cre+ populations of defeated mice (Defeat Cre-: 39.11 ± 6.1 seconds vs. Defeat Cre+: 58.39 ± 9.0 seconds, N=18, 18 mice, respectively, unpaired *t-test p*=0.08). **f.** Cre+ mice, with CA2 activity inhibited, spend significantly less time exhibiting defensive behaviors during defeat when compared to Cre- mice (Defeat Cre-: 44.06 ± 6.3 seconds vs. Defeat Cre+: 24.50 ± 5.2 seconds, N=18, 18 mice, respectively, unpaired *t-test p*=0.02). **g.** Similarly, Cre- mice, on average, spent a significantly higher percent of total submission engaging in more defensive behaviors when compared to Cre+ defeat mice (Defeat Cre-: 52 ± 7.0% vs. Defeat Cre+: 30 ± 4.0%, N=18,18, unpaired *t-test p*=0.01). * *p<*0.05.

### The avoidance phenotype is specific to the acute social defeat experience

Given the understanding that a specific stressor can result in a more generalized stress response [48], we implemented two final behavioral tests using adult, male mice to determine whether or not avoidance of the CD1 aggressor littermate was specific to the aSD stressor. In the first test, the day 1 social defeat stress was replaced with a stressful fear conditioning, consisting of 5 footshocks followed by the day 2 avoidance test and re-exposure to the footshock context (**Supplementary Figure 3a**). As expected, five 0.6 mA footshocks delivered at 30-second intervals resulted in mice displaying increased freezing behavior over the course of the fear conditioning followed by high levels of freezing when the mice were re-exposed to the conditioning context ∼24 hours later (**Supplementary Figure 3b,c**). Furthermore, when shock-conditioned mice were placed in the arena containing a novel CD1 mouse, we observed avoidance levels similar to those of the non-defeated wild type mice and significantly lower than defeated wildtype (**Supplementary Figure 3d**). In a second test, a separate cohort of adult male Amigo2-cre- mice were defeated following our standard aSD protocol and returned to their home cage. Then, 24 hours later, they were placed in the arena containing a novel conspecific (C57BL/6J) mouse and allowed to explore the context. Interestingly, we found that non-defeat control and socially defeated mice spent an equivalent amount of time in the far zone of the arena, opposite the novel mouse (**Supplementary Figure 3f**). From these data we concluded that the socially avoidance phenotype we observe in our defeated population is specific to the aSD experience and not merely a generalized stress response.

### Rostrodorsal CA2 pyramidal neurons preferentially target intermediate CA1 in the caudal hippocampus

Stress responses following exposure to threat or aggression have been shown to affect circuitry within the hippocampus as well as neuronal physiology and signaling in extrahippocampal and limbic brain regions, such as the medial prefrontal cortex (mPFC), anterior cingulate cortex (ACC) and lateral septum [49, 50]. Although CA2 afferents to dorsal lateral septum (dLS) have been shown to underlie aggression towards a conspecific, these pyramidal neurons, like those in dorsal CA1, are not believed to directly project to higher level brain areas like the mPFC. Within the hippocampus, it is the projection neurons located in caudal CA1 (cCA1) that form excitatory connections with such regions [51, 52]. To better understand how CA2 activity may be transmitted to these cortical areas we performed whole cell, patch-clamp experiments in cCA1 of cre+ Amigo2-icreERT2 mice that expressed channelrhodopsin (AAV5-DIO-ChR2-EYFP) in rostrodorsal CA2 (**Figure 4a,b**). This allowed us to record light-evoked excitatory synaptic responses in CA1 pyramidal neurons of coronal slices along the dorsal-intermediate-ventral (DV) axis of the caudal hippocampus originating from rostrodorsal CA2 (**Figure 4c,d**). We observed strong rostrodorsal CA2 (rCA2) axonal labelling in the dorsal and intermediate regions of caudal hippocampal slices (**Figure 4c, Supplementary Figure 4).** Notably, we saw limited EYFP+ labelling in the more ventral region of the cCA1 cell body layer. Consistent with this observation, optical stimulation of rCA2 axons resulted in excitatory synaptic currents in both dorsal and intermediate caudal CA1 cells, with the largest response amplitudes observed in intermediate cCA1 pyramidal neurons, but minimal to no synaptic responses in CA1 cells in the more ventral region of the caudal DV axis (**Figure 4d,e**). The evoked responses we did observe were of a significantly lower amplitude than those recorded at intermediate levels. Interestingly, rCA2 → cCA1 synapses in the dorsal and intermediate areas exhibited paired pulse ratios (PPR) of less than one (*i.e.,* depressing synapses) in response to optical stimulation at a 500 ms interval, with significantly stronger evoked depression occurring in more dorsal CA1 neurons (**Figure 4f,g**). This finding is different than the well-studied CA3 → CA1 Schaffer collateral synapses, which exhibit paired pulse ratios closer to 1 following a 500 ms interval [53, 54].

**Figure 4:**
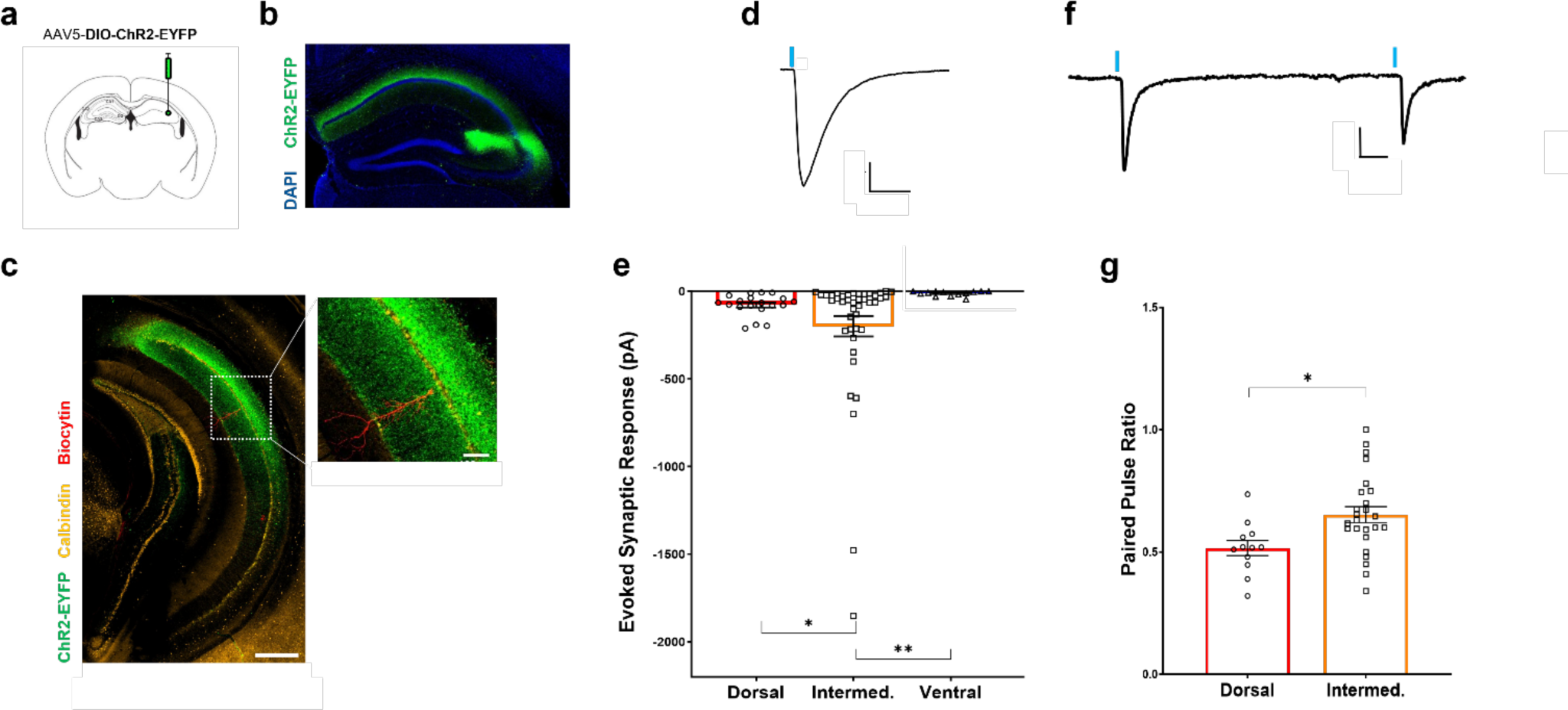
Optical stimulation of CA2 axons evokes responses in caudal hippocampus. **a.** Intracranial injection of 500ul of a Cre recombinase dependent virus containing channelrhodopsin and an EYFP tag were delivered in the rostral CA2 of an Amigo2-icreERT2 mouse. **b.** Image of the resulting channelrhoposin expression pattern in rostral CA2 of in a cre+ Amigo2-icreERT2 mouse (green). **c.** (left) Channelrhodopsin-positive axons of rostral CA2 neurons extended caudally and were the densest in the dorsal and intermediate areas of cCA1 (scale bar = 500 µm). (inset) A biocytin filled pyramidal neuron in dorsal, cCA1 of the hippocampus (red, scale bar = 100 µm). **d.** A 1 ms pulse of laser generated blue light evoked an inward, excitatory current in a cCA1 pyramidal neuron (scale bar = 50 pA x 50 ms). **e.** Light-evoked synaptic responses from CA2 axons were significantly larger in amplitude in neurons found in dorsal and intermediate cCA1 neurons when compared to more ventral locations with the largest response generated in intermediate, cCA1 (Figure 4d**,e**: Dorsal: -77.9 ± 14.0 pA vs. Intermediate: - 199.42 ± 58.0 pA vs. Ventral -12.6 ± 14.4 pA, n=18, 41 and 12 cells, Welch’s ANOVA W=14.67 *p*<0.0001; Dorsal vs. Intermediate unpaired *t-test* with Welch’s correction p=0.048, Dorsal vs. Ventral unpaired *t-test* with Welch’s correction *p*=0.0002, Intermediate vs. Ventral unpaired t-test with Welch’s correction *p*=0.0026). **f.** Two light-evoked pulses, delivered with a 500 ms interval, were used to evoke synaptic response in a cCA1 pyramidal neuron (scale bar = 25 pA x 50 ms). **g.** Both dorsal and intermediate cCA1 neurons showed paired pulse ratios (PPR) of less than 1 indicating these are likely depressing synaptic connections. Dorsal CA1 neurons had a significantly lower PPR than intermediate neurons (Dorsal: 0.52 ± 0.031 vs. Intermediate: 0.65 ± 0.046, n=12 and 25 cells, from 8 and 14 mice, respectively, unpaired *t-test* p=0.012). * *p<*0.05, ** *p*<0.01.

Intrinsic properties of CA1 pyramidal neurons along the dorsal-intermediate-ventral axis of the caudal hippocampus were largely similar with one notable exception: the resting membrane of more ventral CA1 neurons was significantly more depolarized (less negative) when compared to dorsal and intermediate CA1 neurons (**Supplementary Figure 5a**). Action potentials were evoked using 500 ms stepwise current injections and although we observed a trend towards more robust firing in dorsal CA1 cells, we found no significant difference in firing rates between groups at 400 pA current injections (**Supplementary Figure 5b**). The sag ratio elicited by a single hyperpolarizing current injection, a measure of I_h_ current activation, was also similar across neurons from these three regions (**Supplementary Figure 5c**). Finally, it has been proposed that rostrodorsal CA2 neurons preferentially target neighboring calbindin+ CA1 neurons located in the deep, versus superficial, cell layer [25]. In cCA1, we found that the majority of cells that receive synaptic input from rCA2 were, in fact, calbindin negative (**Supplementary Figure 5d**).

### CA2-targeted caudal CA1 pyramidal cells project to corticolimbic brain regions

Our electrophysiological experiments provided evidence that cCA1 neurons in the dorsal and intermediate regions along the DV axis receive excitatory synaptic input from pyramidal neurons in rostrodorsal CA2. As these CA1 neurons have previously been shown to project to extrahippocampal, cortical targets, we next investigated whether cCA1 could be acting as an intermediary in a disynaptic circuit routing CA2 information to higher order limbic brain areas.

We first asked whether activation of CA2 neurons resulted in increased activation of cortical neurons as approximated by cFos staining. To do this, we expressed cre-dependent, excitatory Gq-DREADDs in rCA2 neurons of cre+ and cre-Amigo2-icreERT2 mice via bilateral intracranial injections. After allowing for expression of the Gq-DREADD construct, rCA2 neurons were activated following a single IP injection of CNO (1 mg/kg) and the mice were euthanized 90-minutes later. Brain sections from these mice were immunohistochemically labeled for the immediate early gene (IEG) cFos, one marker for neuronal activity. Although cFos was detectable in cortical areas of cre- control animals, chemogenetic activation of rCA2 drove a significant increase in the number of cFos+ neurons in the ACC of cre+ mice when compared to control mice (**Figure 5a, b**). Similarly, Gq-DREADD activation resulted in a marked increase in the number of cFos+ cells in both the prelimbic and infralimbic subregions of the mPFC of cre+ mice when compared to cre- controls (**Figure 5c,d**). Given our finding that a) rCA2 neurons synaptically activate dorsal and intermediate cCA1 neurons, and b) GqDREADD activation of CA2 results in increased cFos expression in ACC and mPFC, we wondered if dorsal or intermediate cCA1 neurons project to ACC and mPFC. To address this question, we unilaterally infused retrogradely transported AAVs (rAAV-mCherry) into either the prelimbic mPFC or the ACC to label CA1 neurons targeting these areas. In coronal sections of the caudal hippocampus, mCherry+ neurons were found labeled along the dorsal-ventral axis of the CA1 cell body layer and specifically in the dorsal and intermediate regions that receive direct, excitatory input from CA2 (**Figure 5e**). Next, we recorded from acutely prepared caudal hippocampal slices from cre+ Amigo2-icreERT2 mice that expressed ChR2-EYFP in rCA2 neurons and mCherry+ cCA1 neurons with ACC as a known projection target (**Figure 5f**). Optical stimulation of rCA2 axons resulted in excitatory synaptic responses in mCherry+ cCA1 neurons that were significantly larger in amplitude than responses in mCherry-cells that presumably target regions other than the ACC (**Figure 5g**). Taken together, these experiments demonstrate that chemogenetic stimulation of rCA2 is sufficient to drive activation of cells in both the mPFC and the ACC, and that excitatory synaptic information originating in rCA2 pyramidal neurons can be routed to corticolimbic brain areas via cCA1 projection neurons in the dorsal and intermediate regions of the caudal hippocampus. Furthermore, CA2 may preferentially target cCA1 neurons that, in turn, project directly to the ACC.

**Figure 5:**
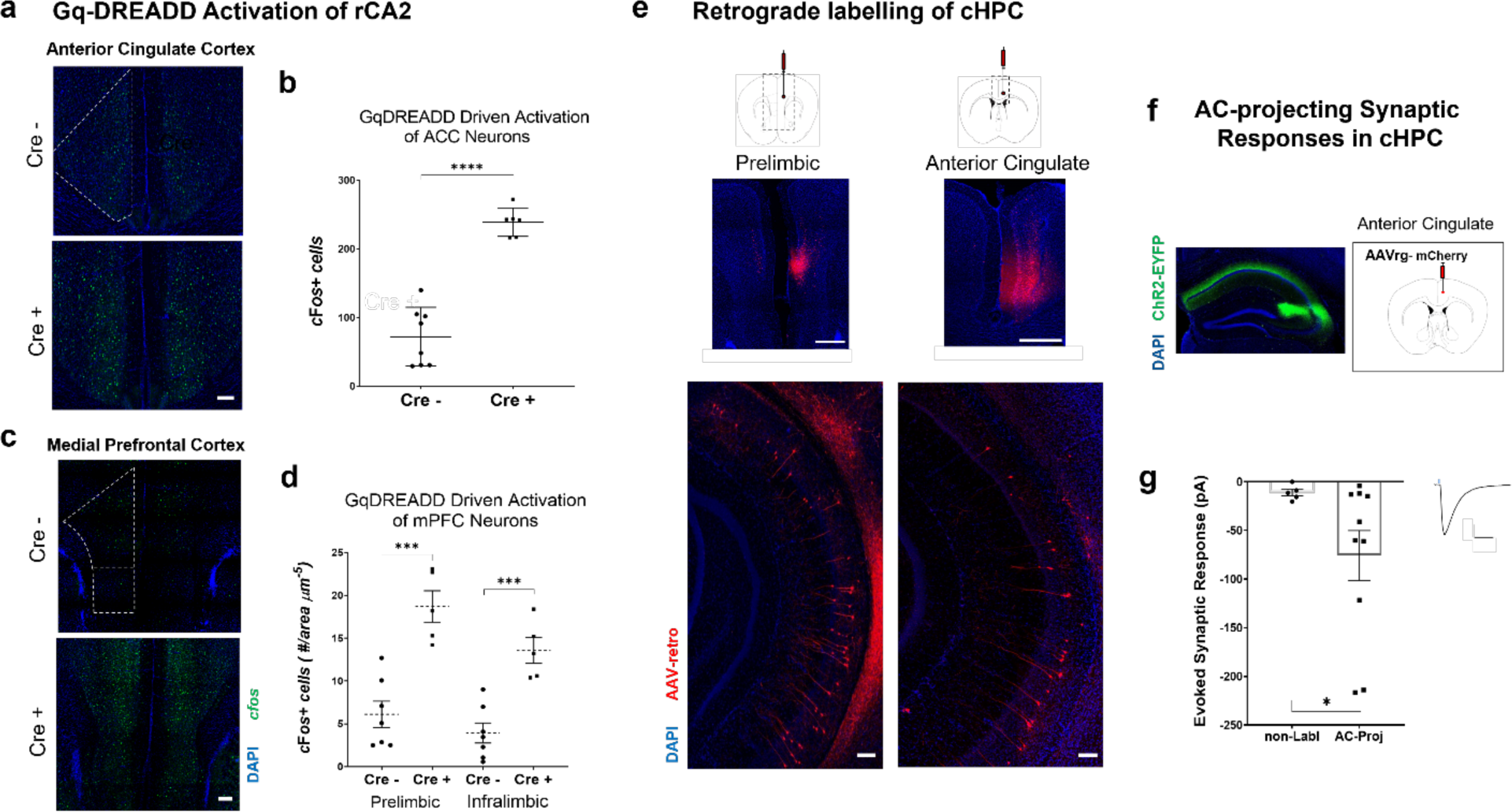
Activation of CA2 neurons engages corticolimbic brain regions, likely through caudal CA1 projection neurons. **a.** Coronal sections from the anterior cingulate cortex of cre- (top) and cre+ Amigo2-icreERT2 (bottom) mice immunohistochemically stained for the IEG cFos (scale bar = 500 µm). **b.** *in vivo* activation of area CA2 via bilaterally expressed Gq-DREADDs causes a significant increase in cFos expression in the ACC of Cre+ mice compared to Cre- mice (Cre-: 72.3 ± 15.1 cFos+ cells vs. Cre+: 239.1 ± 8.5 cFos+ cells, 8 hemispheres from 4 animals and 6 hemispheres from 3 animals, respectively, unpaired *t-test* p<0.0001). **c.** Sections from the prelimbic cortex (mPFC) of cre-Amigo2-icreERT2 (top) and cre+ Amigo2-icreERT2 (bottom) mice immunohistochemically stained for the IEG cFos (dashes outline the more dorsal prelimbic and the more ventral infralimbic cortices, scale bar = 500 µm). **d.** *In vivo* activation of area CA2 via bilaterally expressed Gq-DREADDs drove a significant increase in cFos expression in both prelimbic (Cre-: 6.1 ± 1.5 cFos+ cells/area^-5^ µm vs. Cre+: 18.7 ± 1.8 cFos+ cells/area^-5^ µm, 7 sections from 3 mice and 5 sections from 3 mice, respectively, unpaired *t-test p*=0.0004) and infralimbic (Cre-: 3.9 ± 1.2 cFos+ cells/area^-5^ µm vs. Cre+: 18.7 ± 5.8 cFos+ cells/area^-5^ µm, 7 sections from 3 mice and 5 sections from 3 mice, respectively, unpaired *t-test p*=0.0004) cortex of Cre+ mice compared to Cre- mice. **e.** (left, top) 250-300 µl intracranial injection of AAV-retro-mCherry virus injected into the prelimbic cortex of a wildtype mouse resulted in mCherry expression in neurons along the dorsal-ventral axis of the ipsilateral, cHPC (left, bottom). (right, top) 250-300 µl intracranial injection of AAV-retro-mCherry virus injected into the ACC of a wildtype mouse results in mCherry expression in neurons along the dorsal-ventral axis of the ipsilateral cHPC (right, bottom) (scale bars = 500 µm). **f.** Dual intracranial viral injections resulted in rostral CA2 specific expression of a channelrhodopsin-EYFP (left) and AAV-retro-mCherry labeled neurons in the cHPC neurons that sent axonal projections to the ACC (right). **g.** Rostral CA2 > cCA1 light evoked, excitatory synaptic responses in cHPC neurons that project to the ACC. Responses from neurons that send axonal projections to AC (mCherry+) were significantly larger in amplitude than the synaptic responses from mCherry-neurons that likely project elsewhere (non-labelled cells: -10.28 ± 4.3 pA vs. - 75.8 ± 25.7 pA, n=5 and 10 cells, respectively, *p*=0.034, unpaired *t-test* with Welch’s correction, scale bars = 50 pA x 50 ms). * *p<*0.05, ** *p*<0.01, *** *p*<0.001, **** *p*<0.0001.

### Acute social defeat alters cFos+ expression in frontocortical and hippocampal brain regions during social investigation

In a final set of experiments, we aimed to better understand how the aSD stressor may affect cellular activity in hippocampal and extrahippocampal brain areas during the day 2 avoidance test when subject mice were allowed to investigate an arena containing a littermate of the CD1 aggressor. In this experiment, we examined cFos staining in non-defeated controls and cre- and cre+ defeated mice (given CNO prior to defeat). Forty-five minutes following the avoidance test, animals were euthanized, and coronal brain sections cut and immunostained for cFos. We asked whether cFos would be differentially expressed in the highly avoidant cre+ group, where CA2 activity was inhibited during the defeat (**Figure 6a,c,e**). We found that 24 hours after aSD, the number of cFos+ cells was significantly reduced in the mPFC of cre+ mice when compared to non-defeat controls during social investigation. This effect was largely driven by significantly lower cFos positivity in the prelimbic subregion (**Figure 6b**). Moreover, we observed significantly lower cFos+ neuron counts in highly avoidant cre+ mice in the ACC when compared to non-defeat control mice (**Figure 6b**). Somewhat surprisingly, cFos+ was unaffected in the dorsal lateral septum, a region that receives direct input from rCA2 and modulates aggressive behaviors (**Figure 6b**). In the caudal hippocampus (**Figure 6c**) the number of cFos+ cells in the cCA1 cell body layer was significantly reduced in both cre- and cre+ defeat groups when compared to non-defeat control mice. Interestingly, this effect is mainly driven by a subregion specific loss of cFos positivity in ventral cCA1, a region that our electrophysiological data suggests receives little to no direct synaptic input from rCA2 (**Figure 6d**). These reductions, along with significantly lower cFos expression in caudal CA3 in cre+ mice, result in significantly lower total cFos+ counts in the caudal hippocampus of both cre- and cre+ socially defeated mice. In the rostral hippocampus (**Figure 6e**), a region that underlies social recognition memory [22, 28, 55] and also the region most directly affected by Gi-DREADD mediated inhibition in cre+ mice during defeat, we observed a general trend towards reduced cFos expression in defeated animals. The number of cFos+ neurons was only significantly reduced in the rCA1 of cre-defeated mice when compared to non-defeated control mice (**Figure 6f**). Together these data support the idea that inactivation of CA2 during defeat on day 1 can result in decreased activation of ACC that occurs when animals are investigating another animal on day 2. In contrast, cFos staining in hippocampal areas appeared to best reflect whether an animal had experienced defeat, irrespective of CA2 inactivation status the day before.

**Figure 6:**
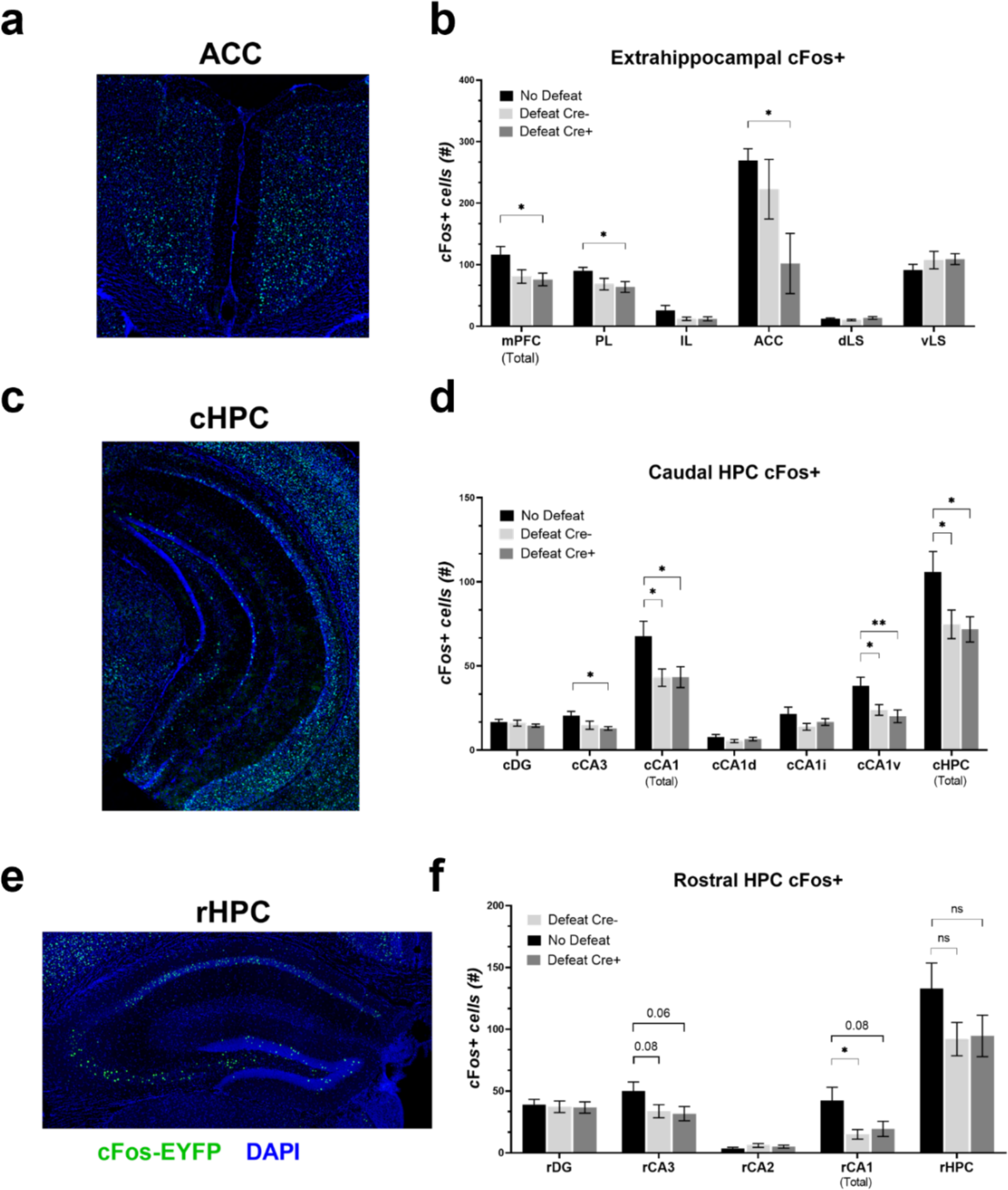
Defeat alters frontocortical and hippocampal neuronal activity in defeated animals during social investigation. **a.** Confocal image of mouse anterior cingulate cortex stained for the immediate early gene (IEG) cFos (green) 45 minutes after the day 2 avoidance test, 20x, scale bar = 100 µm. **b.** Quantification of cFos+ cells in extrahippocampal brain regions active during the investigation of a CD1 mouse 24 hours after aSD. The number of cFos+ neurons in the mPFC, in total, and the prelimbic (PL) subregion were significantly lower in Cre+ mice that had been defeated when compared to No Defeat control animals (mPFC: No Defeat: 116.4 ± 13.3 cells vs. Defeat Cre+: 76.3 ± 10.2, N=5 and 7 mice, respectively, one-way ANOVA F=3.44 *p*=0.059, unpaired *t-test p*=0.035:: PL: No Defeat: 90.3 ± 5.6 cells vs. Defeat Cre+: 64.0 ± 8.7 cells, N=5 and 7 mice, respectively, one-way ANOVA F=2.50 *p*=0.115, unpaired *t-test p*=0.044). Similarly, cFos+ cells were significantly lower in number in the ACC of Cre+ defeated mice compared to No Defeat controls (No Defeat: 269.6 ± 63.0 cells vs. Defeat Cre+: 102.1 ± 48.8 cells, N=11 and 11 mice, respectively, one-way ANOVA F=2.44 *p*=0.95, unpaired *t-test p*=0.048). aSD had no effect on cFos expression in either the dorsal lateral septum (dLS: No Defeat: 12.5 ± 1.3 cFos+ cells vs. Defeat Cre-: 10.4 ± 1.3 cells vs. Defeat Cre+: 13.7 ± 2.2 cells, N=11, 12 and 11 mice, respectively) or ventral lateral septum (vLS: No Defeat: 91.7 ± 9.0 cells vs. Defeat Cre-: 107.7 ± 14.1 cells vs. Defeat Cre+: 109.3 ± 8.8 cells, N=11, 12 and 11 mice) during investigation of a novel CD1 mouse. **c.** Confocal image of mouse cHPC stained for cFOS (green) following the day 2 avoidance test, 20x, scale bar = 500 µm. **d.** cFOS+ neurons counts were significantly lower in cCA3, cCA1 total, ventral cCA1 and the cHPC in total in defeated mice when compared to No Defeat controls (cCA3: one-way ANOVA F=3.32 *p*=0.052, No Defeat: 20.4 ± 2.6 cells vs. Defeat Cre+: 12.9 ± 1.1 cells, N=9 and 10 mice, unpaired *t-test p*=0.014 :: cCA1: one-way ANOVA F=4.14 *p*=0.027, No Defeat: 67.7 ± 9.0 cells vs. Defeat Cre-: 43.1 ± 5.2 cells vs. Defeat Cre+: 43.4 ± 6.2 cells, N=9, 10 and 10 mice; No Defeat vs. Defeat Cre-, unpaired *t-test p*=0.026; No Defeat vs. Defeat Cre+, unpaired *t-test p*=0.037:: cCA1v: one-way ANOVA F=5.59 *p*=0.0095, No Defeat: 38.3 ± 5.0 cells vs. Defeat Cre-: 23.9 ± 3.1 cells vs. Defeat Cre+: 20.1 ± 3.8 cells, N=9, 10 and 10 mice; No Defeat vs. Defeat Cre-, unpaired *t-test p*=0.024; No Defeat vs. Defeat Cre+, unpaired *t-test p*=0.001) :: cHPC total: one-way ANOVA F=3.95 *p*=0.032, No Defeat: 106.0 ± 12.0 cells vs. Defeat Cre-: 74.8 ± 8.5 cells vs. Defeat Cre+: 71.8 ± 7.5 cells, N=9, 10 and 10 mice; No Defeat vs. Defeat Cre-, unpaired *t-test p*=0.046; No Defeat vs. Defeat Cre+, unpaired *t-test p*=0.023). **e.** Confocal image of mouse rostral hippocampus stained for cFOS (green) following the day 2 avoidance test, 20x, scale bar = 100 µm. **f.** Although the number of cFos+ cells trended lower in the rCA3 subregion and for total positive cells, this reduction is only significant when comparing control mice with Cre-defeated animals (rCA3: one-way ANOVA F=2.60 *p*=0.090, No Defeat: 50.0 ± 7.4 cells vs. Defeat Cre-: 33.7 ± 5.3 cells vs. Defeat Cre+: 31.7 ± 5.7 cells, N=11, 13 and 11 mice :: rCA1: one-way ANOVA F=4.16 *p*=0.025, No Defeat: 42.4 ± 10.7 cells vs. Defeat Cre-: 15.0 ± 3.9 cells vs. Defeat Cre+: 19.4 ± 6.1 cells, N=11, 13 and 11 mice; No Defeat vs. Defeat Cre-, unpaired *t-test p*=0.018 :: rHPC total: one-way ANOVA F=1.82 *p*=0.18, No Defeat: 113.0 ± 20.6 cells vs. Defeat Cre-: 92.0 ± 13.5 cells vs. Defeat Cre+: 94.6 ± 16.8 cells, N=11, 13 and 11 mice, respectively). * *p<*0.05.

## Discussion

Here we present evidence that activation of rCA2 neurons is sufficient to drive neuronal activity in corticolimbic targets like the mPFC and ACC. Axons from rCA2 form excitatory synaptic connections with CA1 pyramidal neurons in the dorsal and intermediate regions of the caudal hippocampus (cHPC) and may preferentially target caudal CA1 (cCA1) neurons that project to the ACC. Inhibition of CA2 during an acute social defeat stressor resulted in highly avoidant mice that exhibited significant reductions in defensive behaviors when faced with aggression. Moreover, cFos staining was significantly lower in the rCA2 → cCA1 → ACC circuit in highly avoidant, defeated mice following (the opportunity for) social exploration. Our findings contribute to the increasing understanding of the role that CA2 plays in social behaviors and aggression. Alongside previous work from our lab describing the effects of CA2 inhibition on fear conditioning [32], these data support a role for area CA2 in the processing of stressful stimuli. We also report that the acute social defeat paradigm (aSD) is sufficient to drive social avoidance in a subset of defeated mice, not only one day after the defeat, but also weeks later while the more investigative/resilient mice return to avoidance levels similar to those of non-defeated controls.

Previous studies have identified behavioral and physiological markers that can predict whether an animal would exhibit a more “resilient” or “susceptible” phenotype following chronic social defeat stress [10, 56, 57]. Although much of this work has focused on identifiable variability prior to the stressor, Willmore, et al., observed a defensive, “fighting back” phenotype emerging over the 10-day CSDS paradigm in mice that go on to exhibit resilience following the chronic stress.[13]. Consistent with their finding, our results indicate that inhibition of area CA2 during the aSD can significantly reduce the amount of defensive submission in mice that show higher levels of avoidance (low investigation) when compared to defeated mice with unperturbed CA2 activity (**Figure 3f,g**). If chemogenetic inhibition of CA2 activity is disrupting the CA2 to dorsal lateral septum (dLS) excitatory projections known to positively regulate aggressive behaviors [31], it follows that neuronal activity along this pathway may promote defensive aggression, versus freezing or fleeing, thus leading to a more investigative phenotype following aSD. One question that arises is whether the upright defensive posturing we describe as defensive submission can be considered on the spectrum of aggressive behavior. Our reasoning for categorizing this behavior as submission is two-fold; for one, we only observed this posture in response to aggression and never as active or instigative and two, we did not observe subject mice biting or lashing out towards the aggressor. If defensive submission is akin to aggression, we might expect to see changes in cFos expression in the dLS of defeated mice. Surprisingly, when defeated mice were exposed to a littermate of the CD1 aggressor 24 hours after defeat, we observed no changes in cFos staining in either the dorsal or lateral septum, regardless of whether CA2 was previously active or inhibited (**Figure 6b**, cre- and cre+, respectively). This could be because while active during the defeat, the rCA2 → dLS circuitry may not be reactivated during the day 2 investigation/avoidance test. Interestingly, we also found that CA2 itself has the lowest levels of cFos staining of all the hippocampal areas. However, we know from our *in vivo* and *in vitro* work, CA2 neurons are active during social interactions, but that they have unique molecular features that would be predicted to suppress IEG expression [24, 55, 58, 59]. Future work will help us to better understand the role rostral CA2 may playing in promoting a more defensive response to aggression to support a more investigative/resilient phenotype following an aggressive encounter. CA1 cell bodies span the length of hippocampus along the dorsal-ventral axis in the mouse brain, and cCA1 excitatory projections to the lateral septum, amygdala and cortex have been shown to regulate and modulate anxiety-related behaviors [60–62]. Conversely, the rostrodorsal hippocampus is believed to make few if any afferent connections to higher brain regions that underlie “emotional” behaviors, but more likely send intrahippocampal projections to cCA1 projection neurons that, in turn, project to limbic centers [62–64]. Although much of the literature on hippocampal circuits has focused on the dense concentration of ventral cCA1 projection neurons [28, 30], we found that rCA2 axons preferentially target CA1 neurons in the more dorsal/intermediate regions versus the more ventral region of the cHPC. Consistent with our results, Raam & Sahay demonstrated that optogenetic inhibition of rCA2 afferents terminating in the more dorsal/intermediate region of cHPC was sufficient to disrupt social discrimination during a three-chamber social memory task [26]. Conversely, Meira et al. disrupted social discrimination memory by chemogenetically inhibiting CA2 axons in ventral CA1 (direct injection of CNO). Though we did not observe direct synaptic input from rCA2 onto these more ventral neurons, we did see that reductions in cCA1 cFos expression was primarily found in neurons in vCA1. A possible explanation for the results observed by Meira, et al. is that CNO diffused from the ventral injection location and modulated CA2 axon in more intermediate regions. Moreover, previous work has detailed the reciprocal connectivity both within cCA1 and between cCA1 regions and the subiculum, another important, cortically projecting output region of the hippocampal formation [65]. As we provide evidence that rCA2 sends excitatory signals to dorsal and intermediate cCA1, the afferent connections to either more ventral or subicular projection neurons could further modify hippocampal information before subsequent transmission to corticolimbic regions implicated in stress and anxiety. Future work will be required to better understand not only how the rostral hippocampal information is routed to extrahippocampal, cortical regions, but also how reciprocal connectivity within caudal projection neurons alters or modulates outbound transmission.

Activity in the ACC is modulated by stress. For example, in humans with post-traumatic stress disorder, exposure to emotionally relevant stimuli results in hypoactivity of the ACC when compared to healthy counterparts [66]. In mice, an early-life stressor, such as maternal separation, reduces the activation of ACC pyramidal neurons [67, 68]. Furthermore, repeated social defeat changes ACC IEG level in a manner consistent with the social hierarchy position (Basil, Gross et al. 2018). Caudal hippocampus is known to project to ACC [69], and we found that rostrodorsal CA2 neurons form excitatory synaptic connections onto cCA1 neurons that project to limbic regions such as the mPFC and anterior cingulate cortices. We also found that inhibition of CA2 during aSD (cre+ group, **Figure 2 c, d**) significantly reduces social investigation (*i.e.,* increases “susceptibility”) in defeated mice that also had reduced cFOS+ cells in the ACC. Thus, these findings support the idea that a CA2→cCA1→ACC pathway may at least partially control an animal’s response to social stress acutely, as well as the memory of social stress.

Pyramidal neurons in CA2 express high levels of mineralocorticoid receptors (MRs), but not glucocorticoid receptors (GRs), which prompted this study to examine the role of CA2 in processing information related to a stressful social stimulus [33–35]. Even given the high levels of MRs in CA2, our experiments demonstrate that mice undergoing the fear conditioning (shock) protocol showed minimal avoidance behavior, showing that avoidance is not a generalized response to stress or stress hormones. Rather, the nature of the stressor does seem to be an important factor for later avoidance. Additionally, an animal’s behavior during the defeat (flee vs. defensive behavior) is likely to be a major contributor to later avoidance. Note that the two scenarios, either a) CA2 activity is central for ‘learning’ the identity of an aggressor, or b) CA2 activity is critical for defensive submission, which could be required for the resilient/investigative phenotype, would be exceedingly difficult to disentangle. Also still unknown is whether the resilient/investigative (CA2 activity intact) mice were simply better able to “learn” during the defeat the exact identity of the aggressor (better social discrimination during the defeat), leading them to be less likely to avoid the unfamiliar CD1 mouse. Some level of social discrimination is required for the avoidance behavior, however, in that defeat by a CD1 failed to lead to avoidance of a conspecific. Thus, CA2 activity, be it by modifying behavior in response to an aggressor, and/or by aiding in social discrimination, plays a critical role in the lasting effects of social defeat.

## Acknowledgements

This research was funded by the Intramural Research Program of the National Institute of Environmental Health Sciences, U.S. National Institutes of Health (ES 100221). We thank the expert staff the NIEHS Fluorescence Microscopy and Imaging Center for their help with imaging.

---

**Supplementary Figure 1:**
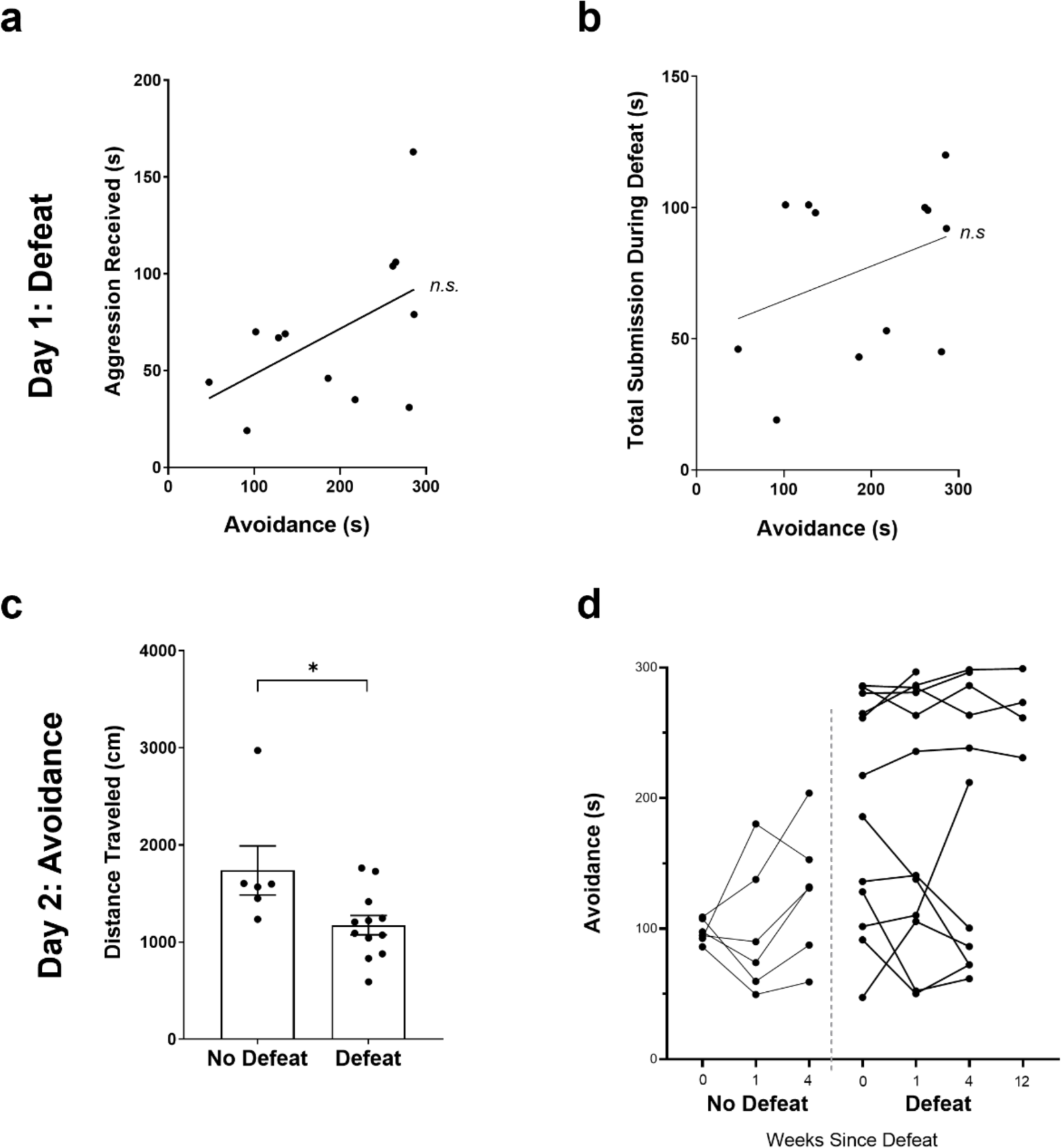
Comparison of subject mouse behavioral metrics during day 1 defeat and day 2 avoidance test. **a.** No significant correlation was observed between the amount of aggression a subject mouse received during the defeat and the amount of avoidance observed 24 hours after defeat (N=12 defeated mice, R^2^=0.2573, *p*>0.05). **b.** No significant correlation was observed between total submissive behavior exhibited by subject mice during defeat and the amount of avoidance observed 24 hours after defeat (N=12 defeated mice, R^2^=0.2727, *p*>0.05). **c.** On average, defeated C57BL/6J mice explored the day 2 arena significantly less than the non-defeated controls during the five-minute avoidance test (No Defeat: 1737.81 ± 253.3 cm vs. Defeat: 1173 ± 99.2 cm, N=6 & 12 mice, respectively, unpaired *t-test p*=0.02). **d.** Highly avoidant, defeated mice (>200 seconds spent in the far zone) remained avoidant for up to one month following a single bout of aSD when exposed to a littermate of the aggressive CD1 when compared to the more investigative, defeated mice (week 0, Defeat-Avoidant: 265.84 ± 10.6 second vs. week 0, Defeat-Investigative: 115.14 ± 19.1 second, N=6 & 6, unpaired *t-test* p<0.0001). * *p<*0.05.

**Supplementary Figure 2:**
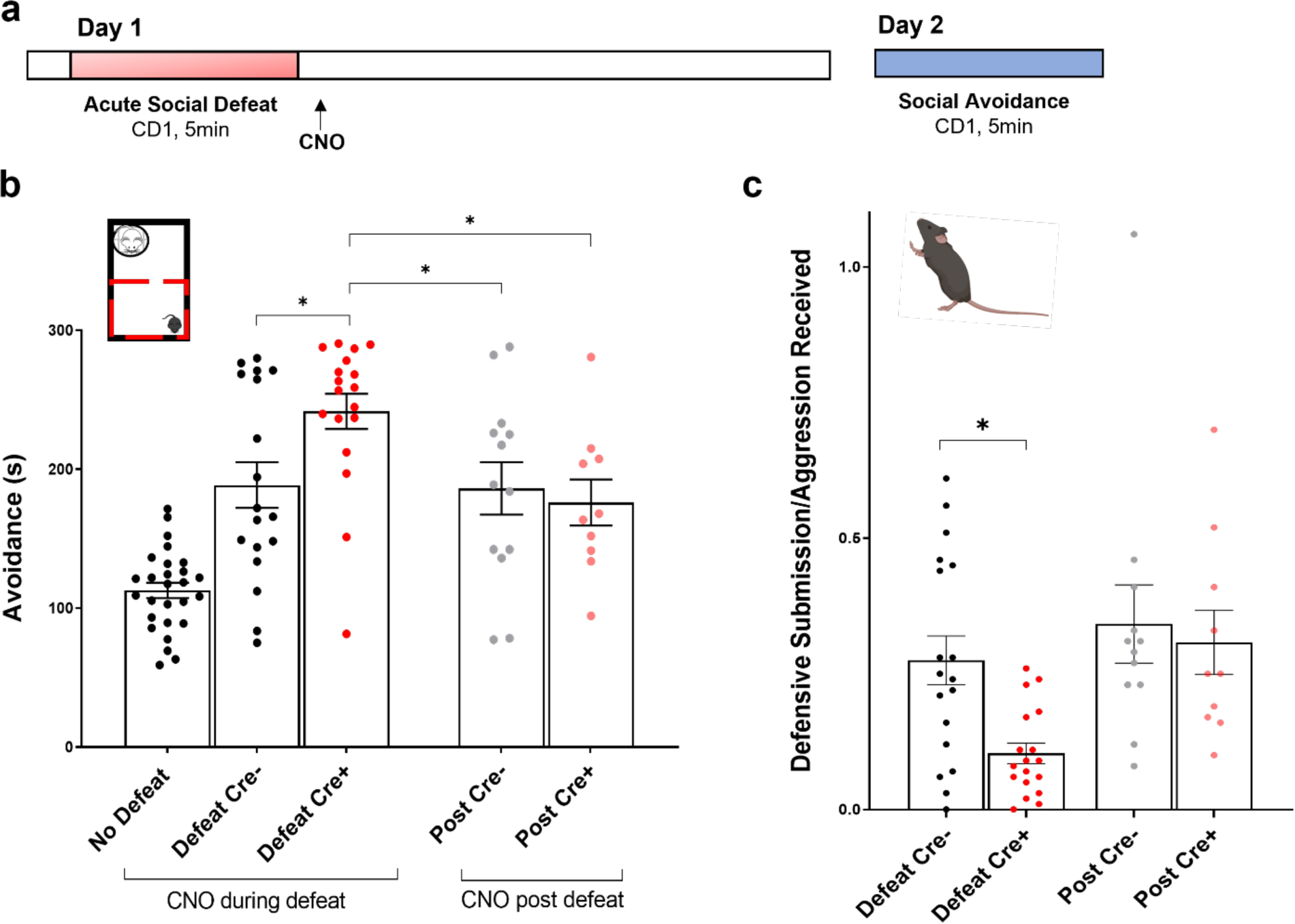
Chemogenetic inactivation of rCA2 after the defeat has no effect on subject avoidance 24 hours post-defeat **a.** Acute social defeat experimental timeline. Five mg/kg of clozapine-N-oxide (CNO) was delivered on day 1, immediately following defeat by a CD1 aggressor mouse via IP injection in Amigo2-icreERT2 male mice (16-19wks of age) that either did (Cre+) or did not (Cre-) express cre-recombinase in area CA2 neurons. Twenty-four hours after defeat, day 2, mice were tested for avoidance/investigation in a novel context containing a littermate of the CD1 aggressor mouse. **b.** Inhibition of area CA2, via the activation of a virally expressed Gi-DREADD construct, immediately following aSD resulted in an avoidance phenotype similar to that of Defeat Cre-animals that received CNO prior to the defeat at 24 hours and significantly lower than Cre+ mice wherein CA2 activity was inhibited during the defeat (data shown are replotted from Figure 2 with additional ‘No Defeat’ animals; Defeat Cre+: 241.64 ± 12.7 seconds vs. Post Cre-: 186.21 ± 18.9 seconds vs. Post Cre+: 176.01 ± 16.6 seconds, N=18, 13, 10, respectively, one-way ANOVA F=16.41 p<0.0001; Defeat Cre+ vs. Post Cre-, Holm-Sidak multiple comparisons test *p*=0.023, Defeat Cre+ vs. Post Cre+, Holm-Sidak *p*=0.016). **c.** Confirmation that the subject defensive behavior during the defeat was similar in mice given CNO after the defeat and cre- mice given prior to defeat. Defensive submission values, normalized to the amount of aggression received, were equivalent to those of defeated Cre- mice that received CNO prior to defeat (Defeat Cre-: 0.28 ± 0.04 vs. Defeat Cre+: 0.10 ± 0.04 vs. Post Cre-: 0.34 ± 0.07 vs. Post Cre+: 0.31 ± 0.06, N=18, 18, 12, 10 mice, respectively, one-way ANOVA F=5.52 *p*=0.0022; Defeat Cre- vs. Defeat Cre+, Holm-Sidak *p*=0.026, Defeat Cre+ vs. Post Cre-, Holm-Sidak *p*=0.004, Defeat Cre+ vs. Post Cre+, Holm-Sidak *p*=0.026). * *p<*0.05, ** *p*<0.01.

**Supplementary Figure 3:**
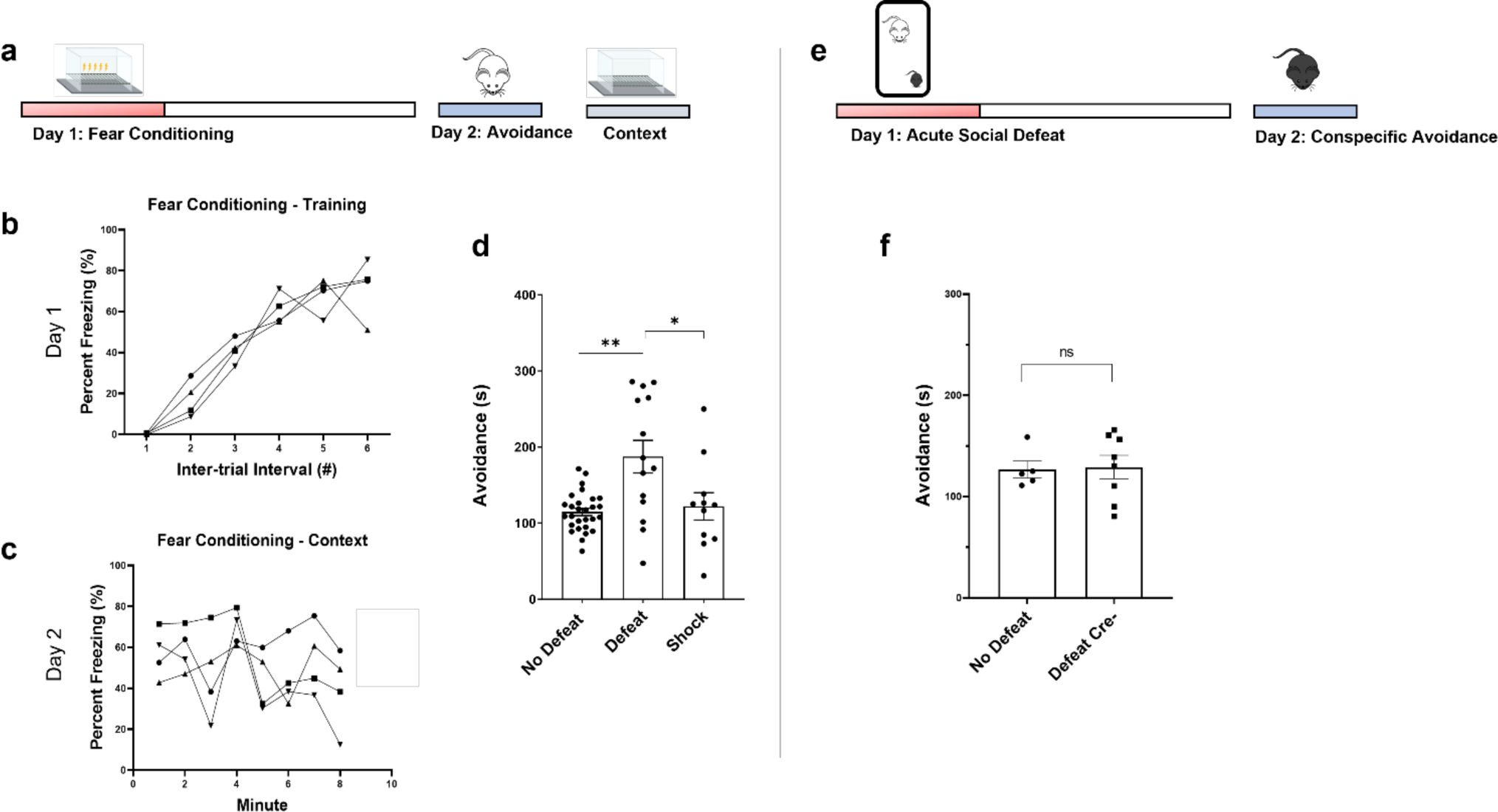
Avoidance is specific to the acute defeat stressor. a. Experimental timeline for a subset of animals in which the aSD stressor was replaced with a five-foot shock, fear conditioning protocol on day 1, followed by a day 2 avoidance test and screening for the acquisition of contextual fear. b. Wildtype mice showed a graded increase percent time spent freezing during the administration of five 0.6 mA footshocks at 30 second intervals (N=4 of 11 total mice tested). c. Twenty-four hours later, mice that underwent fear conditioning continued to exhibit freezing behavior when placed back in the day 1 context (N=4 of 11 total mice tested). d. Twenty-four hours after the administration of a non-social stressor (i.e. footshocks), shock conditioned mice spent a comparable amount of time in the far zone as the no defeat control animals and significantly less time than wild type, defeated mice (No Defeat controls: 114.84 ± 4.8 seconds vs. Defeat: 190.49 ± 25.0 seconds vs. Shock Conditioned: 122.08 ± 18.0 seconds, N=28, 12, 11 mice, respectively one-way ANOVA F=9.47 p=0.0003; No Defeat vs. Defeat, Holm-Sidak multiple comparisons test *p*=0.0003; Defeat vs. Shock Conditioned, Holm-Sidak *p*=0.0063). e. Experimental timeline for a subset of animals that were socially defeated by a CD1 aggressor mouse on day 1 and then allowed to investigate a novel, age matched male conspecific (C57BL/6J) on day 2. f. Socially defeated mice spent an equivalent amount of time in the far zone of a novel arena when allowed to interact with a novel, conspecific 24 hours after being defeated by a CD1 aggressor (No Defeat: 126.70 ± 8.4 seconds spent in the far zone vs. Defeat Cre-: 129.19 ± 11.5, N=5, 8 mice, respectively, unpaired *t-test p*=0.88). * *p<*0.05, ** *p*<0.01.

**Supplementary Figure 4:**
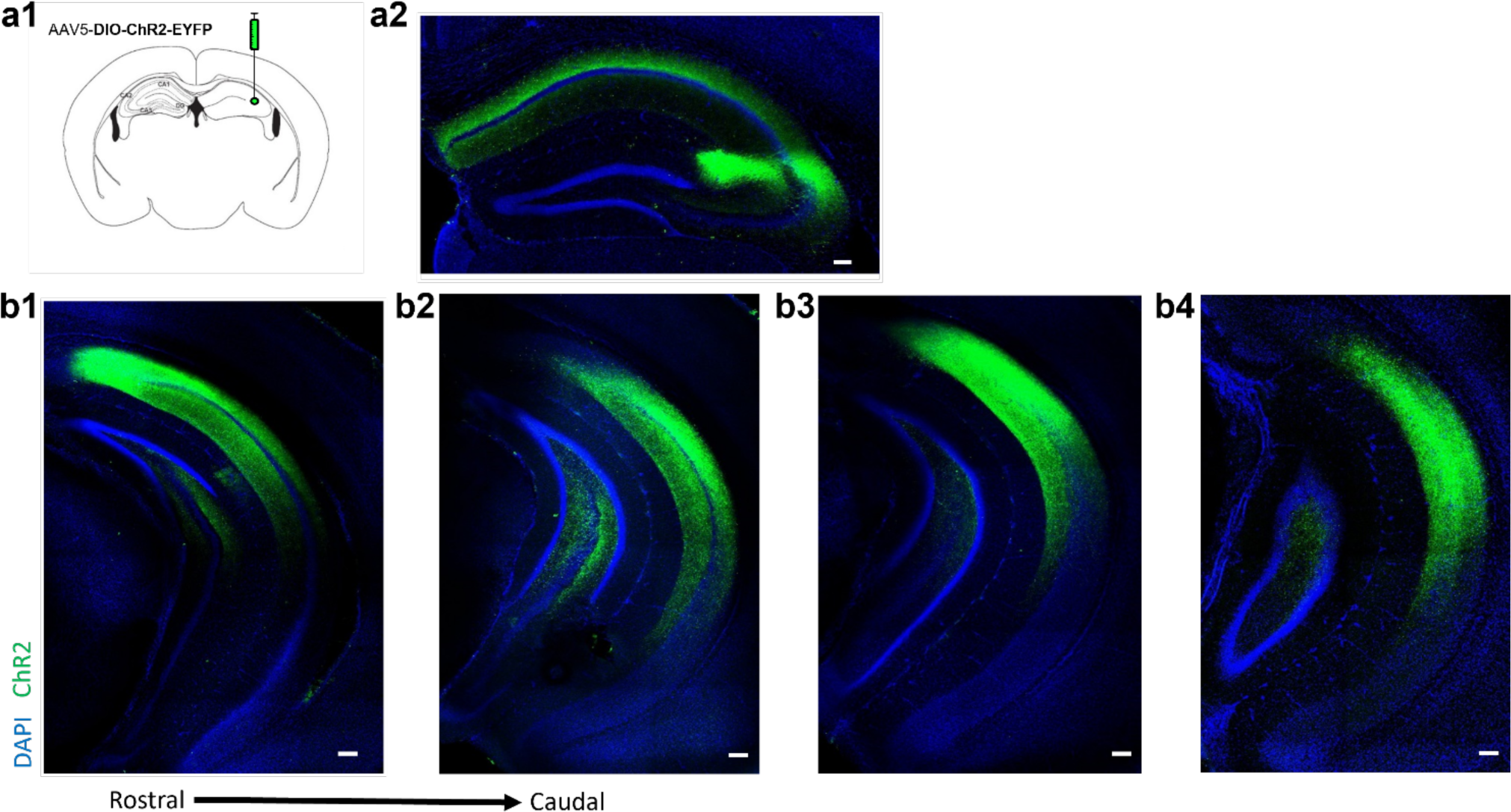
Rostral CA2 axons target the caudal hippocampus in cre+ Amigo2-icreERT2 mice **a1.** Intracranial injection of 500 µl of a cre-recombinase dependent virus containing channelrhodopsin and an EYFP tag were delivered into the rostrodorsal CA2 (rCA2) of an Amigo2-icreERT2 mouse. **a2.** Coronal sections showing the channelrhoposin/EYFP+ (green), cell-type specific expression pattern in rCA2 in an Amigo2-iERT2-cre+ mouse (scale bar = 100 µm). **b1-b4.** EYFP+ axons originating from rCA2 neurons were present in the more dorsal and intermediate extents of caudal hippocampus. The EYFP+ axons were largely absent from the more ventral regions of the cCA1 region (scale bars = 100 µm).

**Supplementary Figure 5:**
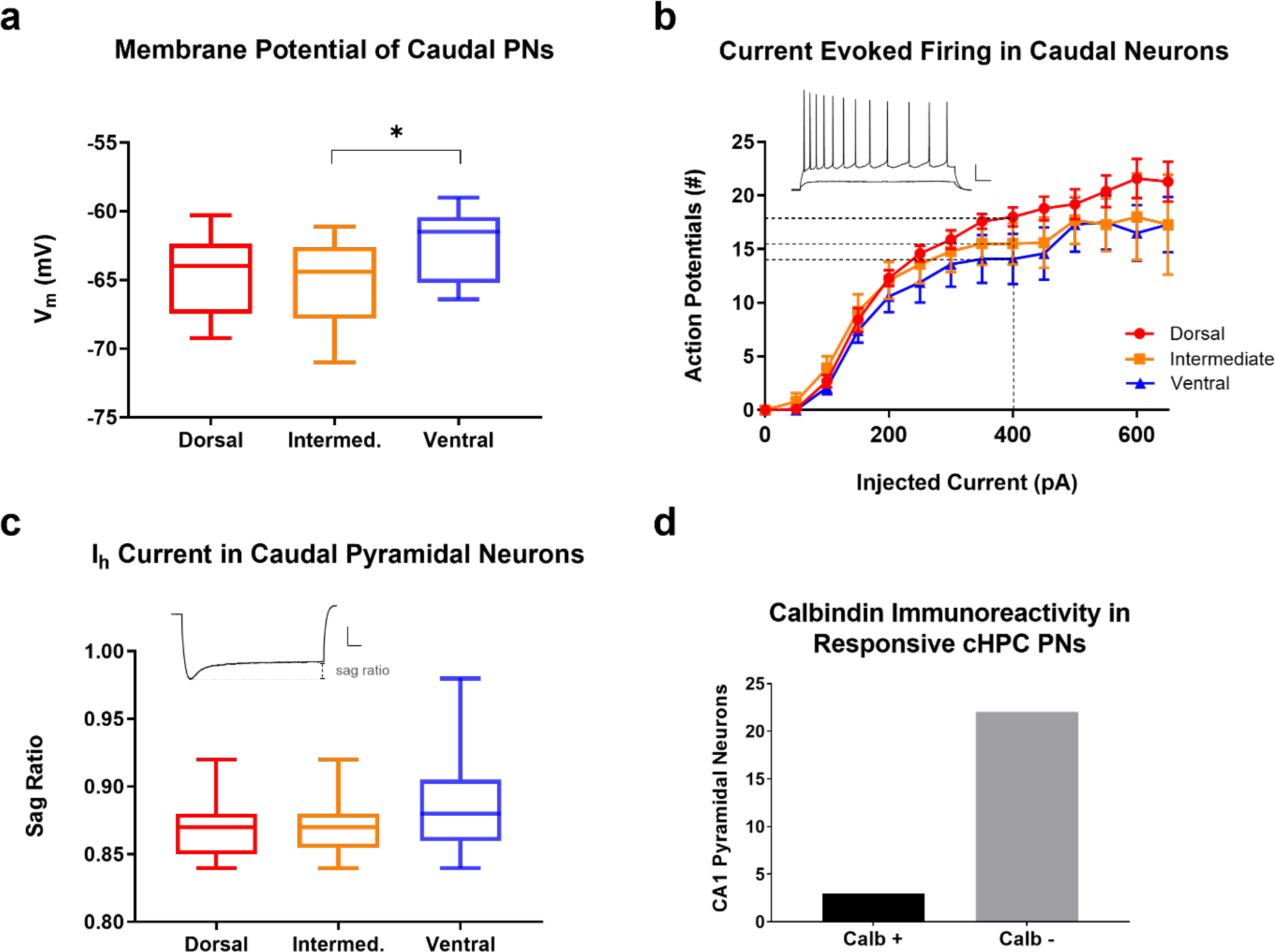
Intrinsic properties of CA1 projection neurons are similar along the D-V axis of the caudal hippocampus (cHPC). **a.** Min/max plots showing the intrinsic membrane potential (V_m_) of cCA1 pyramidal neurons. The average V_m_ of dorsal and intermediate CA1 pyramidal neurons were not significantly different, while intermediate CA1 neurons had a significantly more hyperpolarized membrane potential than ventral cHPC neurons Dorsal: -64.58 ± 0.73 mV vs. Intermediate: -64.95 ± 0.53 mV vs. Ventral: -62.43 ± 0.72 mV, n=16, 31 and 12 cells, from 11, 17, and 7 mice, respectively, one-way ANOVA F=3.468 p=0.038; Dorsal vs. Intermediate, Holm-Sidak *p*=0.67; Dorsal vs. Ventral, Holm-Sidak *p*=0.11; Intermediate vs. Ventral, Holm-Sidak *p*=0.0.035). **b.** cCA1 neurons exhibited similar input/output curves in response to 500 ms steps of depolarizing current (action potentials (APs) elicited with 400 pA current injections; Dorsal: 18.0 ± 0.91 APs vs. Intermediate: 15.5 ± 1.95 APs vs. Ventral: 15.4 ± 2.32 APs, 9 cells from 6 mice, 8 cells from 8 mice, 8 cells from 7 mice, respectively, scale bars = 20 mV x 100 ms). **c.** The voltage sag ratio following a 500 ms hyperpolarizing current injection, a measure of Ih current activation, did not significantly differ in CA1 neurons along the dorsal-intermediate-ventral axis of cHPC (Dorsal: 0.87 ± 0.01 vs. Intermediate: 0.87 ± 0.01 vs. Ventral: 0.89 ± 0.01, n=15, 17 and 9 cells, respectively, scale bars = 20 mV x 100 ms)). **d.** Neurons that received excitatory synaptic input from rostral CA2 are predominately negative for calbindin (Calbindin+ = 3 cells vs. Calbindin- = 22 cells). * *p<*0.05.

